# Overnight circuit remodelling drives juvenile alloparental care

**DOI:** 10.1101/2025.06.13.659524

**Authors:** Bradley B. Jamieson, Maxwell X. Chen, Swang Liang, Lina S. H. El Rasheed, Grace M. K. Chattey, Johannes Kohl

## Abstract

Parental care is critical for the survival of altricial young and is mediated by neural circuits that are well characterised in adult rodents. Although adults can exhibit caregiving even before becoming parents, the developmental origins of this so-called alloparental behaviour remain unclear. Here, we show that alloparental behaviour in mice emerges abruptly between postnatal day (P)14 and 15, independently of prior social experience. This behavioural transition coincides with the onset of pup-specific activity in galanin-expressing medial preoptic area (MPOA^Gal^) neurons, which are essential for parental behaviour in adulthood. Chemogenetic silencing of MPOA^Gal^ neurons abolishes caregiving in juveniles, suggesting that similar circuits control parenting across life stages. Viral trans-synaptic tracing and whole-cell recordings reveal extensive input remodelling of MPOA^Gal^ neurons between P14 and 15, marking a rapid transition from a highly connected, immature network to a sparser, adult-like circuit configuration. We identify microglia as key mediators of this process, as their ablation prevents both synaptic reorganisation and the emergence of alloparenting. Together, these findings uncover a previously unrecognised, microglia-dependent developmental switch that enables caregiving in juveniles through rapid circuit reconfiguration.

## Main

While the neural basis of parental behaviour is extensively well-characterised in adults, the developmental origins and mechanisms by which these circuits become operational remain poorly understood. Parenting is mediated by complex, brain-wide circuits^1–12^, which are shaped by hormonal changes during pregnancy and parturition that strongly promote caregiving in biological parents^13,14^. Yes adult virgin rodents can also display spontaneous parental behaviour^1,13–17^, indicating that core components of parental circuits are functional before reproductive experience. This so-called alloparental behaviour—caregiving provided by non-parents—is typically considered an adult behaviour. However, pup-directed behaviours such as grooming and retrieval to the nest have been observed in weaned juvenile rats^18,19^, prairie voles^20,21^ and mice^22,23^, indicating that parental circuits become operational earlier in life. Yet, the timing and circuit-level mechanisms underlying the emergence of parental behaviour remain unknown. Here we address these questions in mice and show that alloparental behaviour emerges abruptly between postnatal day (P)14 and 15, a process driven by microglia-mediated remodelling of immature circuits into an adult-like configuration. These findings reveal an unexpectedly early onset of caregiving behaviour in mice and provide new insight into how developmental plasticity shapes the maturation of social behaviour circuits.

### Emergence of juvenile alloparenting between P14–15

To address when alloparental behaviour first occurs in mice, we exposed juveniles of both sexes to unfamiliar pups (age P0–2) daily from P14 until P20 (Fig. 1a). P14 was chosen as a start age because eye opening in C57BL/6J mice occurs shortly beforehand, at P13.5 (Extended Data Fig. 3b)^24^. While juveniles showed minimal interactions with pups at P14, we observed a sudden onset of alloparental interactions (pup grooming and retrieval) at P15 (Fig. 1b, c and Extended Data Fig. 1a–c and Supplemental Video 1). Alloparental performance at P15 was comparable to that of adult virgin females (Fig. 1b–c and Extended Data Fig. 1b–c), and individual differences in juvenile performance predicted later alloparental behaviour in adulthood (Fig. 1d). To control for potential effects of repeated pup exposure, a separate cohort was tested at P15 without prior pup exposure. The parental performance of these animals was significantly higher than of animals tested at P14, and comparable to repeatedly exposed P15 mice (Fig. 1e and Extended Data Fig. 1d–g), demonstrating that the onset of juvenile alloparenting does not require prior pup exposure. Both sexes exhibited juvenile alloparenting (Extended Data Fig. 2), and the onset of this behaviour is unlikely to reflect general sensorimotor maturation, because P14 and P15 mice showed no differences in eye opening, auditory startle responses, or locomotor performance (Extended Data Fig. 3b–f). P15 mice showed a preference for interacting with pups over a novel object (toy brick), indicating that their behaviour is directed towards pups rather than driven by novelty (Extended Data Fig. 3g). These findings demonstrate an onset of juvenile alloparental behaviour between P14–15 in mice.

**Fig. 1:**
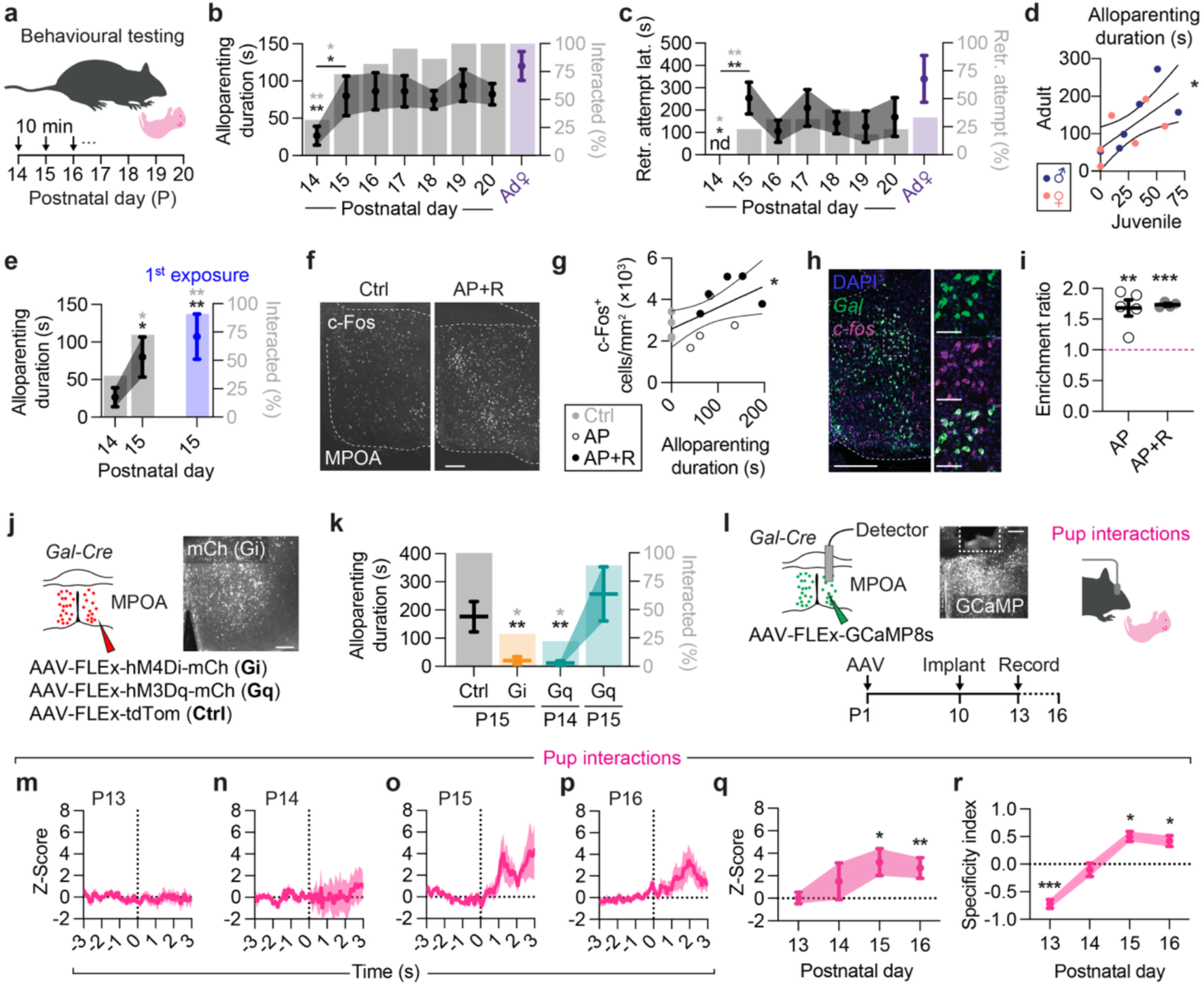
Onset of juvenile alloparental behaviour between P14 and P15 in mice. **a,** Assessing pup-directed behaviour in juvenile mice. **b–c,** Alloparental interactions (**b**) and pup retrieval (retr.) attempts and latency (lat.) (**c**) as a function of age (n = 22 mice). Comparison with alloparental performance of adult virgin females (Ad♀, purple, n = 10 mice) is shown. Right *y* axes show percentage of juveniles that interacted with pups (**b**) or attempted retrieval (**c**). Black and grey asterisks refer to parameters on left and right *y* axes, respectively. Asterisks indicate significant changes between P14 and P15 (horizontal bars), or compared to virgin females; nd, not detected). **d**, Relationship between time spent alloparenting (AP) at P15 (juvenile) compared to re-testing in adulthood (P80, n = 6 males + 6 females, *P* = 0.0146, *R^2^* = 0.4647). **e,** Alloparental performance at P15 without prior pup exposure (n14–15 = 22, n15 first = 11 mice). **f,** Example MPOA sections from non-pup exposed P15 control (Ctrl) and alloparental + retrieval (AP+R) with c-Fos immunostaining. **g,** Relationship between duration of alloparental interactions and density of c-Fos^+^ cells in the MPOA (n = 4 Ctrl, 4 AP, 5 AP+R, *P* = 0.0248, *R^2^* = 0.4105). **h,** *Gal* and *c-fos* transcripts in the MPOA visualised via smFISH. Scale bars, 500 µm (inset 100 µm). **i,** Enrichment of *c-fos* (see Methods) in MPOA^Gal^ neurons relative to non-exposed controls (n = 5, 4 mice). Asterisks indicate significant changes compared to controls (dashed line). **j,** Chemogenetic inhibition (Gi) or activation (Gq) of MPOA^Gal^ neurons; example MPOA section shown. Scale bar, 200 µm. **k,** Alloparenting during MPOA^Gal^ manipulation (n = 4 Ctrl, 7 Gi, 7 Gq). Asterisks indicate significant differences from tdTomato controls in alloparenting duration (black) or percent of mice interacting with pup (grey). **l,** MPOA^Gal^ fibre photometry: GCaMP8s expression, fibre placement, and experimental timeline. Scale bar, 250 µm. **m–p,** Averaged MPOA^Gal^ activity during pup interactions (n = 10). Dashed lines mark interaction onset). **q**, Peak Z-scored activity during pup interactions; asterisks indicate significant differences from baseline. **r,** Tuning Specificity Index of Z-scored MPOA^Gal^ activity during pup interactions, calculated relative to responses to biological mother, unfamiliar dam, adult virgin female, mouse milk and siblings (n = 10, see Methods). Positive values indicate stronger responses to pups than to other stimuli (i.e., pup-specific tuning), values near zero reflect non-selective responses, and negative values indicate weaker responses to pups than to other stimuli. Asterisks indicate significant differences from zero). Statistics: Kruskal-Wallis test with Dunn’s *post hoc* and Holm correction (**b, e**), Kaplan-Meier survival analysis log-rank test with Benjamini-Hochberg FDR (**c**), Fisher’s exact test with Benjamini-Hochberg FDR (**b, c, e, k**), linear regression (**d, g**), one-way ANOVA with *post hoc* Dunnet’s test (**i**), unpaired *t*-test with *post hoc* Tukey’s HSD (**k**), *U* test with Dunn’s *post hoc* and Holm correction (**q**), Wilcoxon signed-rank test with Benjamini-Hochberg FDR (**r**). Data are mean ± s.e.m. (error bars or shading).

### Juvenile alloparenting recruits neurons critical for adult parental behaviour

The hypothalamus, septal, and amygdaloid nuclei contain several neuronal populations essential for pup-directed behaviour in adults^3,7,25^. To address which brain areas within these regions are engaged during juvenile alloparenting we analysed expression of the immediate early gene *c-fos*—a proxy for neuronal activation—following alloparental interactions involving pup grooming alone (AP) or pup grooming with retrieval (AP+R) (Extended Data Fig. 4). Among the activated regions, the medial preoptic area (MPOA)—a central node in parenting circuits^26–28^—showed a positive correlation between c-Fos^+^ cell number and duration of pup-directed behaviours (Fig. 1f, g). Single-molecule fluorescent *in situ* hybridisation (smFISH) revealed that *c-fos* expression was enriched in galanin-expressing (MPOA^Gal^) neurons—a population essential for adult caregiving—after juvenile alloparental behaviour, regardless of whether retrieval occurred (Fig. 1h, i)^1,2^. Core components of the adult parenting circuit are therefore also recruited during juvenile alloparental behaviour. To probe the functional contributions of MPOA^Gal^ neurons to juvenile alloparenting—and to determine whether the onset of caregiving could be accelerated or suppressed—we chemogenetically manipulated their activity (Fig. 1j). Silencing MPOA^Gal^ neurons significantly impaired alloparental behaviour, indicating that they are required for its expression (Fig. 1k). By contrast, chemogenetic activation at P15 produced a modest but non-significant enhancement of caregiving and activation at P14 failed to elicit alloparental responses (Fig. 1k). These observations suggest that neurons critical for adult parental behaviour also contribute to juvenile alloparenting, but that maturation of additional downstream circuit elements is required for MPOA^Gal^ neurons to drive this behaviour before P15.

To assess potential activity changes in MPOA^Gal^ neurons during the onset of juvenile alloparenting, we performed *in vivo* fibre photometry recordings in juveniles that were exposed to pups and other social stimuli (Fig. 1l and Extended Data Fig. 5). MPOA^Gal^ neurons exhibited significantly increased activity during pup interactions from P15 onwards (Fig. 1m–q). In contrast, these neurons were selectively activated during interactions with adult females prior to P15 (Extended Data Fig. 5a, c, d, f, h, i). Quantification of tuning specificity (see Methods) confirmed that robust pup-specific responses first appeared at P15 (Fig. 1r). MPOA^Gal^ neurons thus become activated during pup interactions between P14–15, coinciding with the onset of alloparental behaviour.

### Remodelling of MPOA^Gal^ inputs during the onset of juvenile alloparenting

Juvenile alloparenting circuits may become functional between P14–15 due to changes in MPOA^Gal^ neuron number, remodelling of their downstream projections or synaptic inputs, and maturation of their intrinsic biophysical properties. We therefore examined whether alloparental onset at P15 was accompanied by circuit-level changes. The density of MPOA^Gal^ neurons did not significantly differ between P14 and P15 and was ∼3-fold lower compared to adults (Extended Data Fig. 6). We mapped MPOA^Gal^ axonal projections by injecting Cre-dependent, adeno-associated viruses (AAVs) encoding the fluorophore tdTomato and the presynaptic marker synaptophysin conjugated to GFP (SynGFP) into the MPOA of P14, P15 and adult virgin female *Gal-Cre* animals (Extended Data Fig. 7a, b). The density of tdTomato-labelled fibres and SynGFP-positive, putative synapses was similar between P14, P15 and adults (Extended Data Fig. 7c–h), indicating that MPOA^Gal^ axonal projections do not undergo major changes after P14.

Next, we addressed putative developmental changes in MPOA^Gal^ synaptic inputs using rabies virus-mediated retrograde trans-synaptic tracing (Fig. 2a and Extended Data Fig. 8a)^29^. Because rabies virus requires ∼4 days to effectively label MPOA^Gal^ presynaptic inputs^1^, we could not assess changes in input patterns at 1-day resolution. Mice therefore received rabies injections at either P10 (‘pre-P14’) or P15 (‘post-P14’)—and were sacrificed 4 days later, at P14 or P19, respectively—to capture MPOA^Gal^ synaptic inputs before or after the putative P14–15 remodelling event (Fig. 2b, c). We compared the resulting labelling patterns to those in adult virgin females (Fig. 2d). Pre-P14, MPOA^Gal^ neurons received extensive, brain-wide inputs, with a high ratio of presynaptic-to-starter neurons (convergence index, CI = 60.1 ± 11.7) (Fig. 2b, e and Extended Data Fig. 8b–d). In contrast, inputs were much sparser when rabies was injected post-P14 (CI = 15.8 ± 5.9), resembling the adult pattern (CI = 6.6 ± 3.9) (Fig. 2c–d and Extended Data Fig. 8c–d). To quantify differences in age-specific MPOA^Gal^ input patterns, we performed Principal Component Analysis (PCA)-based dimensionality reduction on these input distributions and calculated Euclidean distances between age groups in principal component (PC) space (Fig. 2g). Distances between pre-P14 and post-P14, as well as between pre-P14 and adults, were significantly greater than shuffled controls, whereas post-P14 and adult profiles were similar (Fig. 2h). Almost half (29/62) of MPOA^Gal^ input areas exhibited reduced labelling after P14, with pronounced sparsening observed in the midbrain, pallidum, hippocampus, cortex and olfactory areas (Fig. 2f and Extended Data Fig. 8c). These findings suggest that a remodelling event after P14 profoundly alters the input landscape of MPOA^Gal^ neurons.

**Fig. 2:**
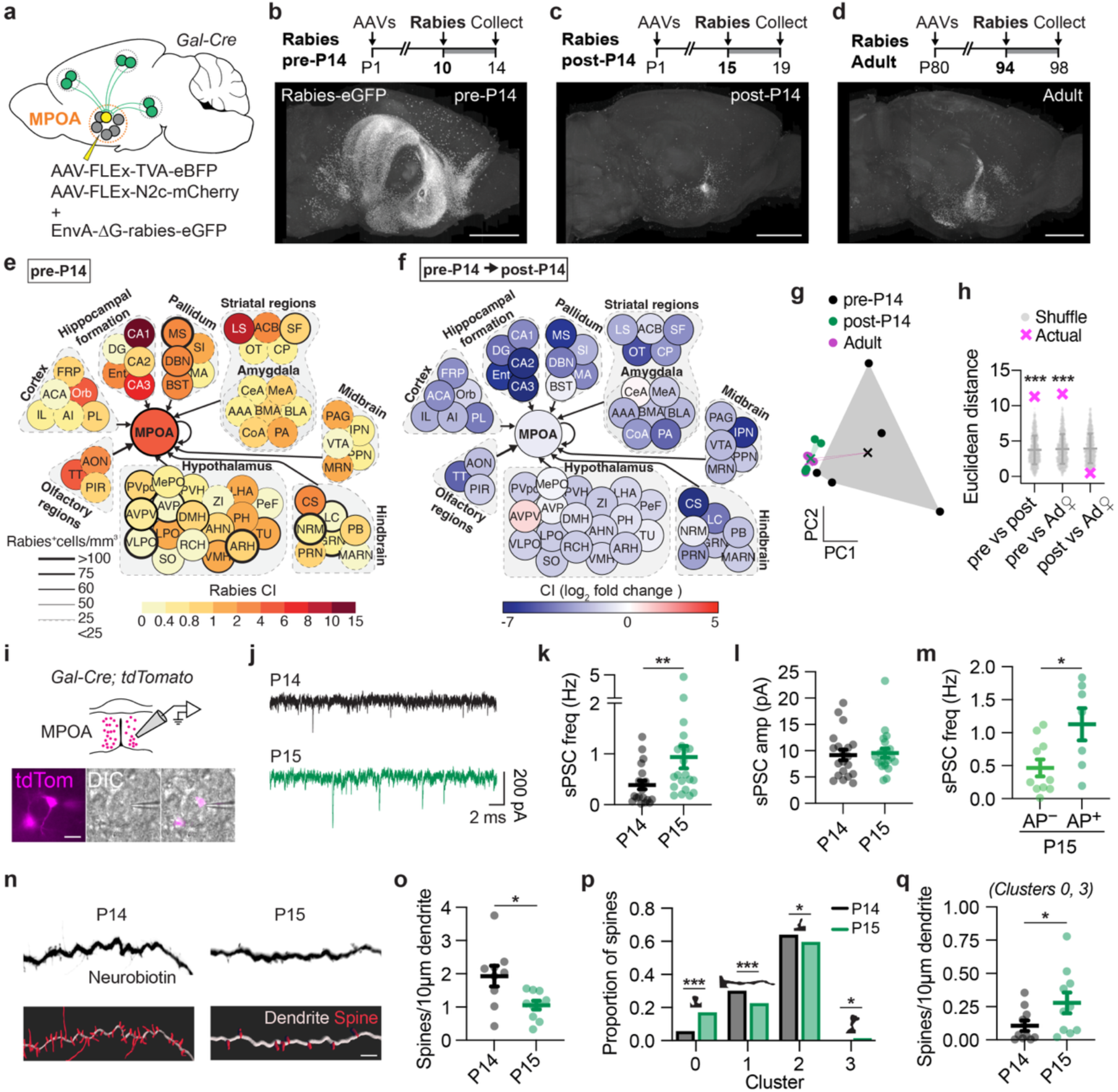
Remodelling of MPOA^Gal^ inputs coincides with the onset of juvenile alloparenting. **a,** Monosynaptic retrograde rabies tracing from MPOA^Gal^ neurons. **b–d**, Timeline of surgeries (top) and sagittal projections of brains imaged via serial section two-photon tomography with rabies-labelled inputs pre-P14 (**b**), post-P14 (**c**) and in adult virgin female (**d**) mice. Scale bars, 2 mm. **e,** Input density and average convergence index (CI) across regions at pre-P14 (n = 5 mice). **f**, Change in CI from pre- to post-P14 (n = 5, 5 mice). **g,** Input distributions in PC space. Each dot is an animal. **h,** Euclidean distances between age-specific input profiles at pre-P14, post-P14 and in adults (n = 5, 5, 4) compared to shuffle (mean ± SD). **i,** Whole-cell recordings from MPOA^Gal^ neurons at P14 and P15. **j,** Example voltage clamp traces with sPSCs. **k–l,** sPSC frequency (**k**) and sPSC amplitude (inward currents, **l**) at P14 and P15 (20, 22 neurons from n = 5, 5 mice). **m,** sPSC frequency from non-alloparental (AP^−^) and alloparental (AP^+^) P15 mice (10, 7 neurons from n = 2, 1 mice). **n,** Neurobiotin-filled dendrites from P14 and P15 MPOA^Gal^ neurons (top) with 3D reconstructions and spines (bottom). Scale bars, 5 µm. **o,** Spine density (9, 10 neurons from n = 6, 5 mice). **p,** Change in prevalence of dendritic spine clusters from P14 to P15; examples shown. **q,** Density of spines in clusters 0 and 3 (see **p;** 9, 10 neurons from n = 6, 5 mice). Statistics: permutation test vs shuffle (n = 1,000) (**h**), *U* test with *post hoc* Dunn’s test with Holm correction (**k– m, p, q**), Fisher’s exact test with Benjamini-Hochberg FDR (**p**). Data are mean ± s.e.m.

We next asked whether the increased number of presynaptic inputs at P14 could be validated using other measures of connectivity. The largest age-related difference in presynaptic labelling was observed in the hippocampus, which provided 25.1 ± 7.0% of inputs to MPOA^Gal^ neurons pre-P14, but less than 5% post-P14 (4.5 ± 1.9) and in adults (4.7 ± 1.5), corresponding to a ∼30-fold decrease in CI between pre-P14 and adults. To verify this decrease in connectivity after P14, we injected AAVs expressing tdTomato and SynGFP into the hippocampus and subsequently examined labelling in the MPOA (Extended Data Fig. 9a). Although tdTomato^+^ fibre density in the MPOA was similar between P14 and adults, SynGFP^+^ puncta were approximately three times more abundant at P14 (Extended Data Fig. 9b–d), indicating denser synaptic contacts from the hippocampus at this age. Pronounced input remodelling thus stabilises MPOA^Gal^ afferent connectivity into an adult-like pattern shortly after P14.

We previously found that estradiol and progesterone remodel MPOA^Gal^ neurons during pregnancy, thereby promoting parental behaviour^14^. Because similar proportions of MPOA^Gal^ neurons express receptors for these hormones in juveniles and in adults (Extended Data Fig. 10a–f), and given the transient activation of the hypothalamic-pituitary-gonadal axis in juvenile mice (‘minipuberty’)^30–32^, we tested whether ovarian hormones also contribute to MPOA^Gal^ remodelling in juveniles. Monosynaptic rabies tracing from adult females ovariectomised at P7 resulted in adult input patterns similar to controls (Extended Data Fig. 10g). MPOA^Gal^ input remodelling is thus not mediated by ovarian hormones in juveniles, consistent with the lack of a sexual dimorphism in juvenile alloparenting (Extended Data Fig. 2).

To determine whether altered input patterns are reflected in individual MPOA^Gal^ neurons— and whether the biophysical properties of these neurons change during the onset of alloparenting—we performed whole-cell patch clamp recordings in brain slices (Fig. 2i). MPOA^Gal^ did not show significant changes in intrinsic properties between P14–15 (Extended Data Fig. 11). However, we observed a selective increase in the frequency of spontaneous postsynaptic currents (sPSCs) at P15—with no change in sPSC amplitude (Fig. 2j–l)— suggesting a presynaptic mechanism. Importantly, sPSC frequency was selectively increased in P15 animals that exhibited alloparental behaviour (Fig. 2m), linking enhanced MPOA^Gal^ inputs with the expression of juvenile alloparenting. Morphological reconstructions of recorded neurons (see Methods) revealed an unexpected overall reduction in dendritic spine density at P15 (Fig. 2n, o). Unsupervised clustering of spine shapes showed that this was primarily due to a loss of long, thin filopodia spines—typically regarded as immature and silent (Fig. 2p)^33^. Concurrently, shorter, mushroom-like spines—which are more likely to contain mature, active synapses^33^—were more prevalent (Fig. 2p, q). Thus, despite a net loss of spines, synaptic connectivity is enhanced through pruning of immature synaptic contacts and enrichment of functionally mature ones, likely accounting for the increased sPSC frequency at P15. In parallel, MPOA^Gal^ neurons exhibited reduced axonal length and branching at P15, suggesting additional fine-scale maturation of MPOA^Gal^ projections (Extended Data Fig. 12a–b, m–p). Together, these data confirm that MPOA^Gal^ synaptic inputs are reorganised between P14–15, coinciding with the onset of alloparental behaviour.

### Microglia-mediated remodelling of MPOA^Gal^ inputs drives onset of juvenile alloparenting

We next asked which mechanisms drive this rapid input remodelling. We focussed on microglia, due to their well-established role in synaptic pruning during development^34–36^. While the density of microglia in the MPOA remained stable between P14–15 (Fig. 3b), their morphology underwent striking morphological changes: microglia at P15 appeared smaller, rounder and less complex (Fig. 3a, c–e and Extended Data Fig. 13a, g–i), consistent with a shift toward and activated, phagocytotic state. In contrast, astrocyte density and morphology were unchanged, and oligodendrocyte-lineage cells showed reduced density and increased cell size, likely reflecting ongoing maturation (Extended Data Fig. 13b–d, j–r)^37^. Unsupervised clustering confirmed a ∼4-fold increase in the proportion of ameboid microglia at P15, as well as a corresponding decrease in highly branched morphologies (Fig. 3f). Similar changes occurred across the hypothalamus (Extended Data Fig. 14a), but not in other brain regions such as the motor cortex or striatum (Extended Data Fig. 15). Notably, these morphological alterations were already detectable by P14.5, suggesting they are established within a 12-hour window (Extended Data Fig. 14b–k). Next, to test whether MPOA microglia are involved in synaptic remodelling of parenting circuits, we examined their engulfment activity. We expressed tdTomato in Gal^+^ neurons and performed immunostainings against microglial and lysosomal markers (Fig. 3g), allowing us to assess whether microglia internalise material from MPOA^Gal^ neurons. At P15, neuron-microglia distance was reduced and a larger fraction of microglial lysosomes contained tdTomato (Fig. 3h, i), despite similar overall numbers of lysosome-containing microglia (Fig. 3j). These findings suggest that MPOA microglia phagocytose cellular material from Gal^+^ neurons between P14–15.

**Fig. 3:**
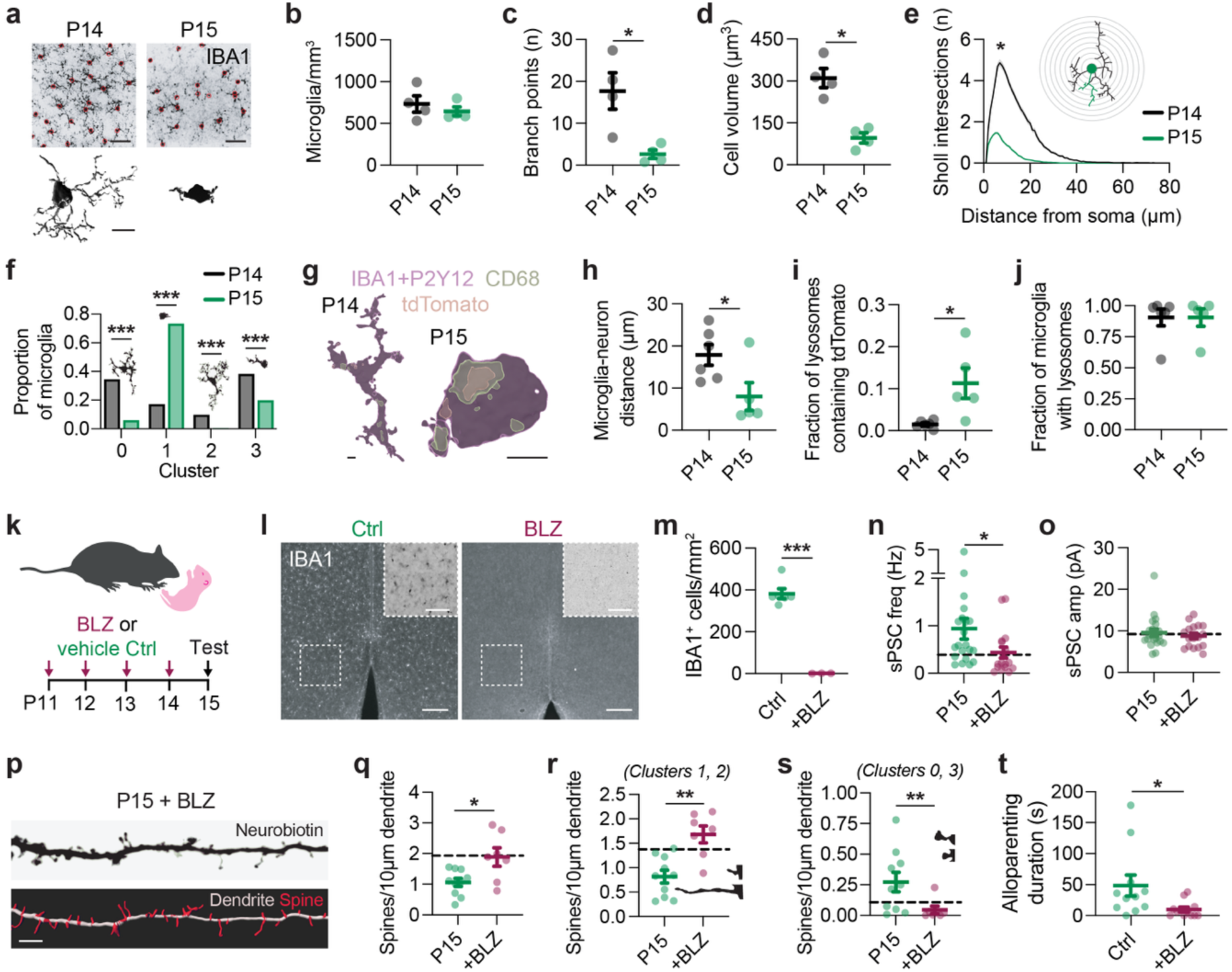
Microglia-mediated remodelling of MPOA^Gal^ inputs drives onset of juvenile alloparental behaviour. **a,** Confocal images of IBA1-immunostained (top) and reconstructed (bottom) MPOA microglia at P14 and P15. Red circles indicate cell bodies. Scale bars, 20 µm (top), 10 µm (bottom). **b–e,** Morphological analysis of MPOA microglia at P14 and P15 (n = 4, 4 mice): cell density (**b)**, branch points (**c**), total cell volume (**d**), Sholl analysis (**e,** example skeletons and 2 µm radius zone shown). **f,** Change in prevalence of morphological clusters from P14 to P15; example microglia morphologies shown**. g,** Reconstructed microglia containing tdTomato^+^ debris in CD68^+^ lysosomes. Scale bars, 5 µm. **h–j,** Microglia proximity to tdTomato^+^ neurons (**h**), fraction of lysosomes containing tdTomato (**i**), and fraction of microglia with lysosomes (**j,** n = 6, 5 mice). **k,** Microglia ablation via i.p. injection of BLZ945. **l,** IBA1 staining in MPOA following vehicle (Ctrl) or BLZ treatment. Insets are colour-inverted for visibility. Scale bars, 200 µm (insets 100 µm). **m,** IBA1^+^ cell density in MPOA of control and BLZ-treated mice (n = 6, 3 mice). **n–o,** sPSC frequency (**n**) and amplitude (**o**) of MPOA^Gal^ neurons in P15 control and BLZ-treated mice (22, 18 neurons from n = 5, 4 mice). **p,** Neurobiotin-filled dendrites (top) and spine reconstructions (bottom) from a P15 MPOA^Gal^ neuron after BLZ-treatment. Scale bar, 5 µm. **q–s,** Overall spine density (**q**), and density of spine clusters with immature (**r**) or mature (**s**) morphology (see Fig. 2p); example spines shown (10, 7 neurons from n = 5, 5 mice). **t,** Alloparental behaviour in control- and BLZ-treated P15 mice (n = 11, 12). Dashed lines in **n–o**, **q–s** indicate average P14 values. Statistics: *U* test with *post hoc* Dunn’s test and Holm correction (**b–e, h–j, m–o, q–t**), Fisher’s exact test with Benjamini-Hochberg FDR (**f**). Data are mean ± s.e.m. (error bars or shading).

Finally, to test whether microglia play an instructive role in remodelling MPOA^Gal^ synaptic inputs, we ablated them in juvenile mice using the CSF1R-inhibitor BLZ945 (Fig. 3k–m) and assessed the resulting biophysical and morphological properties of MPOA^Gal^ neurons. Microglial ablation reduced the frequency—but not amplitude—of synaptic inputs at P15, thereby mimicking a P14-like state (Fig. 3n, o). This manipulation also increased overall dendritic spine density to P14-like levels (Fig. 3p–q), primarily through an increase in spines with immature morphology (Fig. 3r). In contrast, we observed a decrease in density of spines with mature, mushroom-like morphology (Fig. 3s). Microglial ablation therefore disrupts the synaptic pruning of inputs onto MPOA^Gal^ neurons between P14–15. Apart from reduced action potential halfwidth and evoked spike latency, microglia ablation did not affect intrinsic neuronal properties (Extended Data Fig. 16). Strikingly, BLZ-treated juveniles exhibited reduced pup-directed alloparental behaviour compared to controls (Fig. 3t), despite showing otherwise normal pup contact latencies and locomotion (Extended Data Fig. 17). Microglia are therefore critical for the structural and functional maturation of MPOA^Gal^ inputs, and their absence impairs the synaptic remodelling required for the onset of juvenile alloparenting.

## Discussion

Our study identifies a critical developmental switch for the onset of caregiving behaviour in juvenile mice. We show that alloparental interactions start at P15—much earlier than previously reported (Fig. 1a–c and Extended Data Fig. 1a–c)^38^. Using immediate early gene mapping, *in vivo* imaging and chemogenetic manipulations, we demonstrate that MPOA^Gal^ neurons are critical for this early-life behaviour (Fig. 1f–r, Extended Data Fig. 4, and Extended Data Fig. 5), mirroring their role in adult parenting^1,2^.

While the intrinsic properties of MPOA^Gal^ neurons remain stable between P14–15 (Extended Data Fig. 11), their synaptic landscape undergoes pronounced remodelling: whole-cell recordings and single-neuron reconstructions reveal an overnight increase in synaptic inputs, paralleled by dendritic spine maturation (Fig. 2i–q). This remodelling is also apparent from monosynaptic rabies tracing, but surprisingly, the number of neurons presynaptic to MPOA^Gal^ neurons decrease after P14 (Fig. 2a–h and Extended Data Fig. 8). While it remains largely unknown what determines the efficiency of rabies spread between neurons^39^, our results suggest that it scales with the total number of synaptic contacts—irrespective of synapse maturity. The origin(s) and roles of restructured inputs into MPOA^Gal^ neurons will be a subject for future investigations, but the onset of MPOA^Gal^ population activity during pup interactions at P15 (Fig. 1m–q) suggests that they convey pup-related sensory information. Up until P14, MPOA^Gal^ neurons are broadly responsive during interactions with potential caregivers—biological mothers, unfamiliar dams, and virgin females—but these responses diminish by P15 (Extended Data Fig. 5), when MPOA^Gal^ neurons become selectively tuned to pup interactions (Fig. 1r). This shift in tuning from caregiver to pup cues could result from the replacement of upstream inputs or a change in tuning at earlier stages in the circuit (e.g., in peripheral sensory neurons or thalamic relays). Before P14, MPOA^Gal^ neurons receive the largest fraction of their inputs from the hippocampus, but this connection is pruned within days (Fig. 2b–c, f). The functional relevance of these transient inputs remains uncertain, but they may reflect collateral innervation by hippocampal axons en route to other hypothalamic targets^40^. Supporting this, hippocampal axon density within the MPOA remains stable between P14 and adulthood, while the number of putative synaptic contacts declines (Extended Data Fig. 9). An intriguing alternative is that these transient inputs functionally contribute to juvenile alloparenting.

In contrast to the pronounced input remodelling, MPOA^Gal^ outputs remain largely stable during this period even though subtle changes in axonal fibre density (Extended Data Fig. 7g–h) and morphology (Extended Data Fig. 12n–p) hint at additional downstream plasticity. This is also supported by the finding that chemogenetic activation of MPOA^Gal^ neurons at P14 is not sufficient to elicit alloparental behaviour (Fig. 1k).

The onset of alloparental behaviour does not seem to be driven by social learning as it is not accelerated by early pup exposure (Fig. 1e and Extended Data Fig. 1d–g). Moreover, it does not reflect a general maturation process: we found no major sensorimotor differences between P14–15 (Extended Data Fig. 3), and the associated microglial changes appear confined to the hypothalamus (Extended Data Fig. 14a and Extended Data Fig. 15). Our observations thus point to a circuit- and behaviour-specific reorganisation, although this remains to be tested in other hypothalamic cell types. Recent transcriptomic and chromatin accessibility profiling has revealed diverse, cell type-specific postnatal developmental trajectories in the preoptic area^41^, supporting the notion of discrete windows of remodelling in different MPOA populations.

We identify microglia as key mediators of parental circuit remodelling in juveniles, consistent with the role of these cells in activity-dependent synaptic pruning during critical periods of neural development^42–44^. Microglial ablation maintains a P14-like MPOA^Gal^ neuronal phenotype at P15—including overall elevated spine densities and reduced sPSC frequency (Fig. 3n, q)—and impairs the emergence of alloparental behaviour (Fig. 3s). Although microglia ablation has been reported to elevate auditory thresholds^45^, BLZ-treated juveniles in our study contacted pups with latencies and at rates comparable to controls, and exhibited no other overt behavioural impairments (Extended Data Fig. 17). These findings confirm that microglia play an instructive, rather than a merely supportive, role in MPOA^Gal^ circuit reorganisation. Since the density of mature dendritic spines is increased at P15 (Fig. 3r), microglia seem to promote spine maturation in addition to pruning immature spines and stabilising existing mature ones (Fig. 4). Local dendritic calcium elevations are thought to attract microglia via several possible cellular signals that promote either spine maturation or removal^43,44,46–50^, but how MPOA^Gal^ dendritic spines are specifically targeted remains to be determined.

**Fig. 4:**
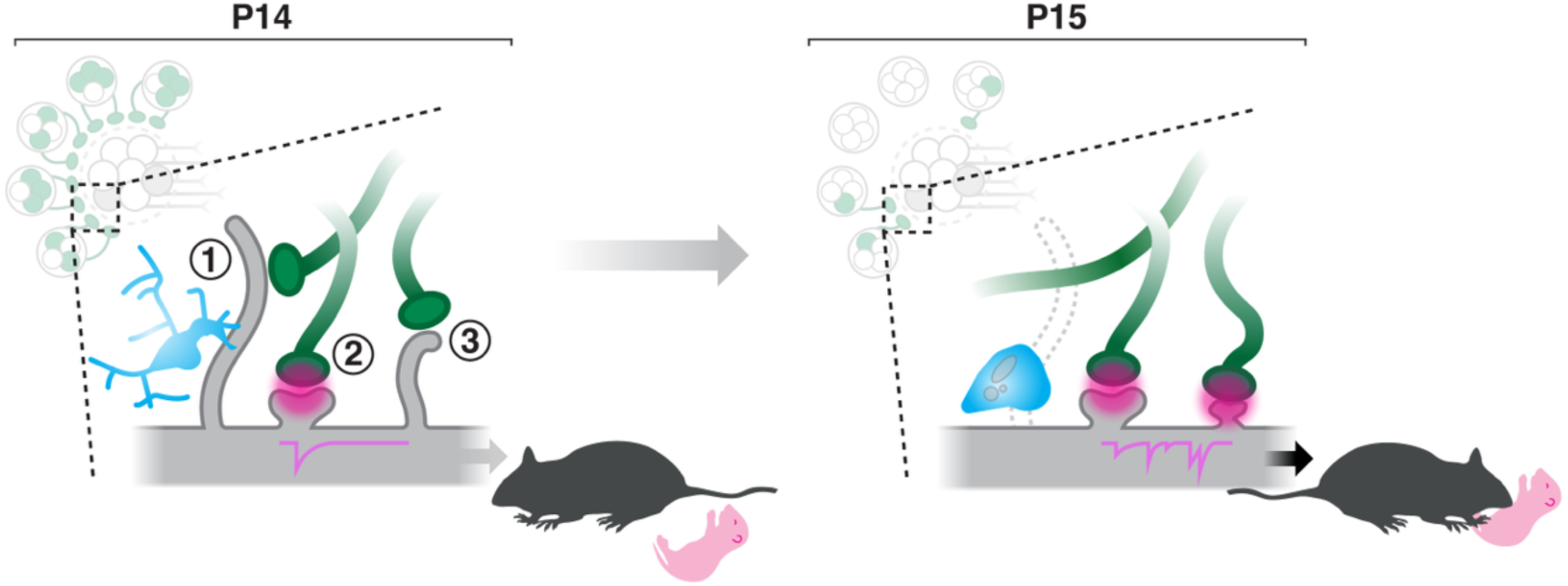
Model of microglia-mediated circuit remodelling underlying juvenile alloparental behaviour. At P14, parenting-promoting MPOA^Gal^ neurons receive extensive, brain-wide inputs but their dendritic spines are predominantly immature, forming weak or transient synapses with axons conveying pup-relevant information. Consequently, MPOA^Gal^ neurons exhibit low postsynaptic activity and are not activated by pup cues. At this stage, MPOA microglia display a complex, ramified morphology consistent with a surveying state. Between P14–15, these microglia adopt an ameboid morphology and remodel MPOA^Gal^ synaptic inputs—likely by pruning immature spines (1), stabilising mature ones (2), and facilitating spine maturation (3). This remodelling results in overall fewer but more functional synaptic inputs, enabling MPOA^Gal^ neurons to drive alloparental behaviour.

Importantly, juvenile alloparenting is not sexually dimorphic (Extended Data Fig. 2) and the associated circuit remodelling does not depend on ovarian hormones (Extended Data Fig. 10), in contrast to what has been observed during pregnancy^51^. Microglial ablation enhances pup-directed behaviours in adult virgin female rats via an unknown mechanism^52^, raising the possibility that microglia-mediated circuit reorganisation may also contribute to behavioural transitions later in life.

What is the ethological relevance of juvenile alloparenting? In the wild, mice live in breeding collectives^53,54^, where auxiliary caregiving by juvenile and adult virgin females can increase offspring survival^55^. Indeed, we find that the extent of juvenile alloparental behaviour predicts adult caregiving performance (Fig. 1d), suggesting continuity in parental circuit function across life stages. While the adult virgin males in our study were spontaneously parental— likely reflecting housing conditions (see Methods)—virgin males are often infanticidal^56,57^. This implies that additional remodelling occurs in male parenting circuits after P15, shifting their pup-directed behaviour from caregiving to aggression.

Improved functional access to developing neural circuits in rodents has begun to reveal how specific neuronal populations support life-stage-specific behaviours^58–62^. For example, agouti-related peptide (AgRP) neurons, which regulate hunger in adults, modulate social attachment in neonates and juveniles^58,59^, while somatostatin neurons in the zona incerta transition from regulating neonatal social behaviour to fear and anxiety in adults^61^. Together with our findings, these studies highlight that instinctive behaviour circuits remain malleable throughout development to support age-appropriate behavioural repertoires. We identify microglia-mediated reconfiguration of synaptic inputs as a mechanism underlying such transitions, providing a framework for investigating how behavioural changes are coordinated across the lifespan.

## Extended Data Figures

**Extended Data Fig. 1:**
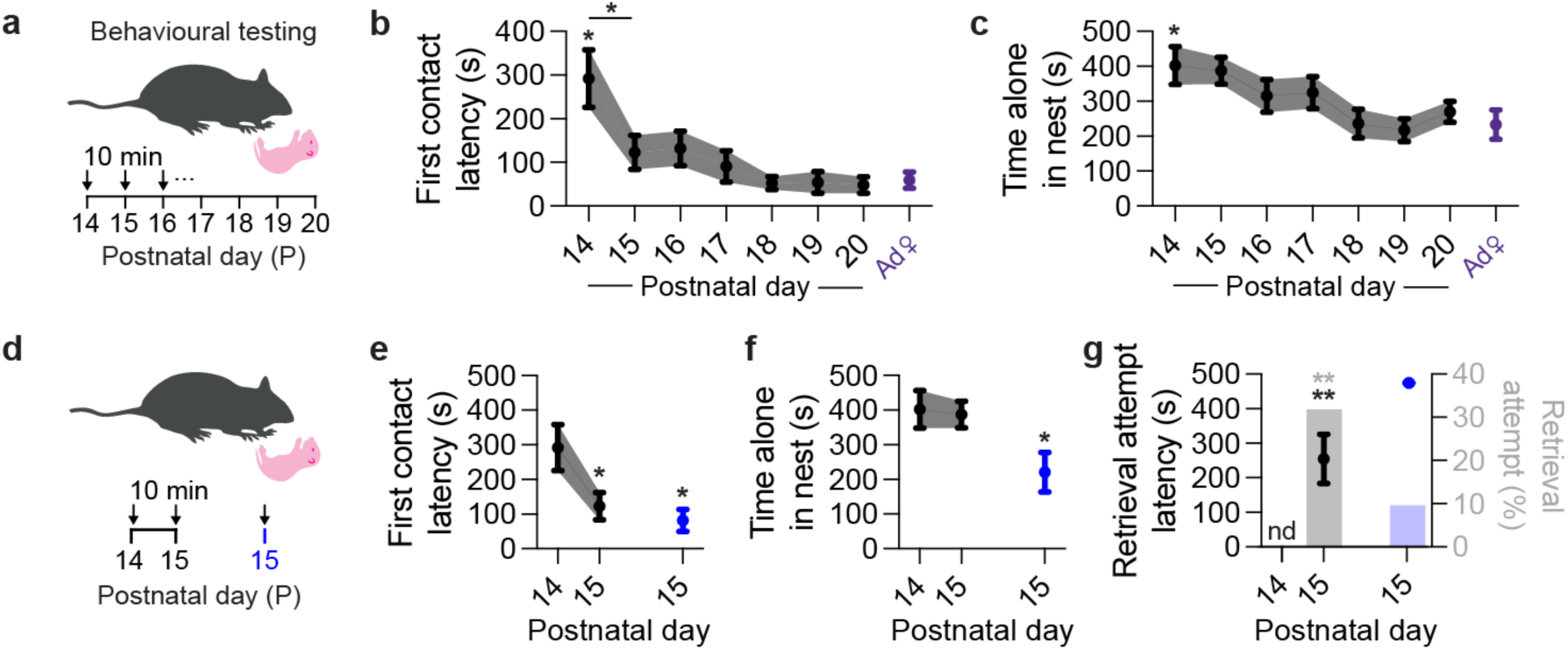
Characterisation of juvenile alloparental behaviour. **a,** Assessing pup-directed behaviour in juvenile mice. **b–c,** Pup contact latency (**b**) and time spent in nest (**c**) as a function of age (n = 22 mice). Values for adult females (Ad♀, purple) are from a separate cohort of adult virgin females (n = 10 mice). Asterisks indicate significant changes between P14 and P15 (horizontal bars) or compared to virgin females. **d,** Comparing pup interactions between juveniles that were repeatedly tested (P14, P15) or first tested at P15 without prior pup exposure. **e–g,** Pup contact latency (**e**), time spent in nest (**f**), and retrieval attempt latency and percentage of mice attempting retrievals (**g**) in repeatedly tested P15 mice and P15 mice without prior pup exposure (n = 22, 11). Asterisks indicate significant differences compared to P14. Statistics: Kruskal-Wallis test followed by Dunn’s *post hoc* test with Holm correction for multiple comparisons (**b, c, e, f**), Kaplan-Meier survival analysis log-rank test with *post hoc* Benjamini-Hochberg FDR correction (**g**), Fisher’s exact test with Benjamini-Hochberg FDR correction (**g**). Data are mean ± s.e.m. (error bars or shading).

**Extended Data Fig. 2:**
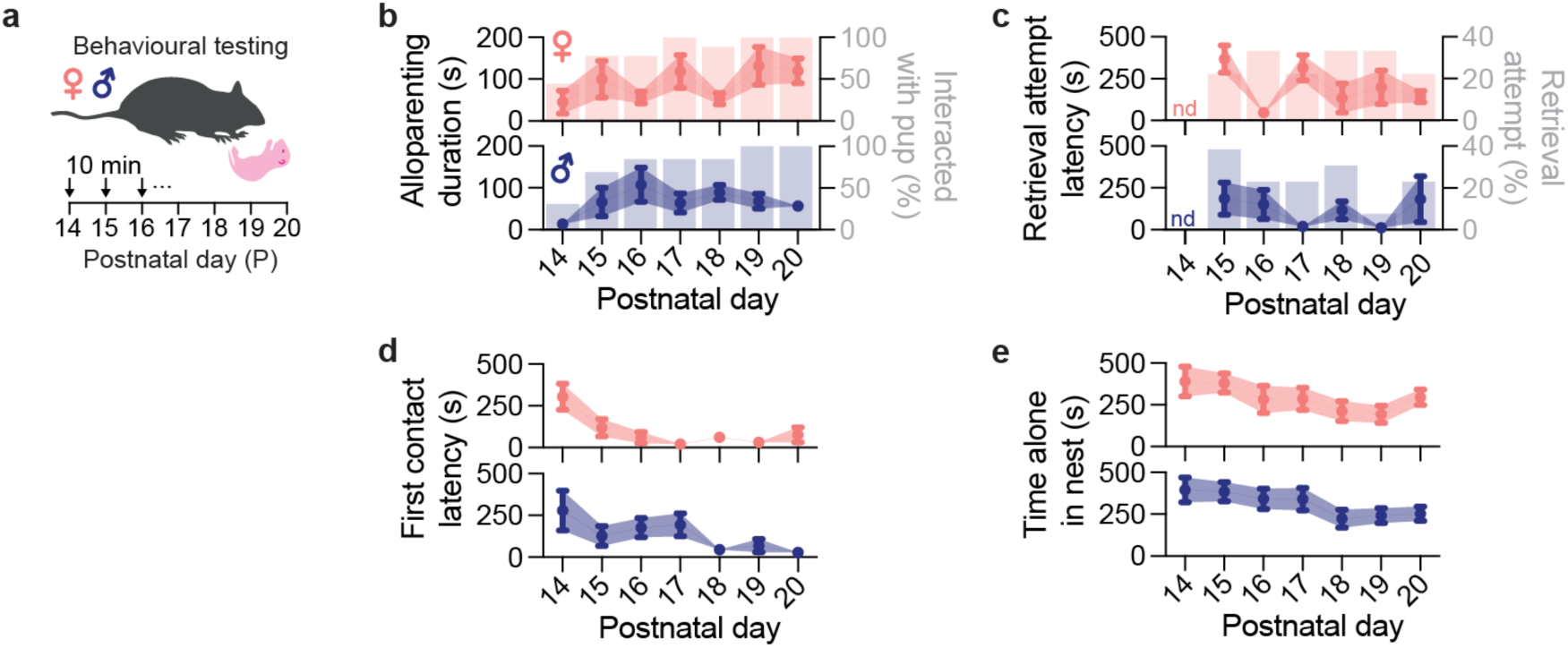
Development of juvenile alloparental behaviour in females and males. **a,** Assessing pup-directed behaviour in female and male juvenile mice. **b–e,** Duration of alloparental interactions and percentage of mice interacting with pups (**b**), retrieval attempt latency and percentage of mice attempting retrievals (**c**), pup contact latency (**d**), and time spent in nest (**e**) of female and male juveniles (n = 9, 13). Statistics: *U* tests with *post hoc* Dunn’s test with Holm correction (**b–e**) and Fisher’s exact test with Benjamini-Hochberg FDR correction (**b, c**). Data are mean ± s.e.m. (error bars or shading).

**Extended Data Fig. 3:**
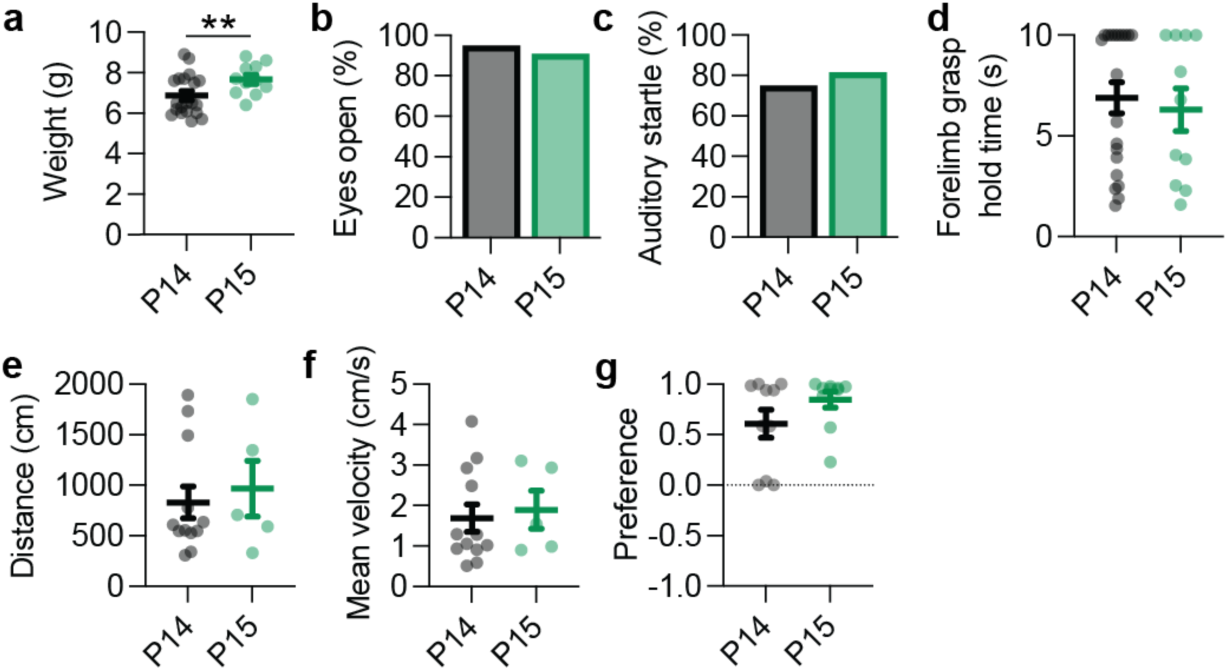
Comparison of sensorimotor function between P14 and P15. **a–f,** Mouse weight (**a**), percentage of mice with fully open eyes (**b**), percentage of mice displaying auditory startle reflex in response to a clapping stimulus (**c**), forelimb grasp hold time in 10 s assay (**d**), distance moved in 10 min open field assay (**e**), and mean velocity in 10 min open field assay (**f,** n = 20, 11 mice). **g**, Novel object preference index between pup and toy brick (1 = preference for pup, −1 = preference for toy brick). Statistics: *U* test with *post hoc* Dunn’s test with Holm correction (**a, e– g**), Fisher’s exact test (**b, c**), Kaplan-Meier survival analysis log-rank test (**d**). Data are mean ± s.e.m.

**Extended Data Fig. 4:**
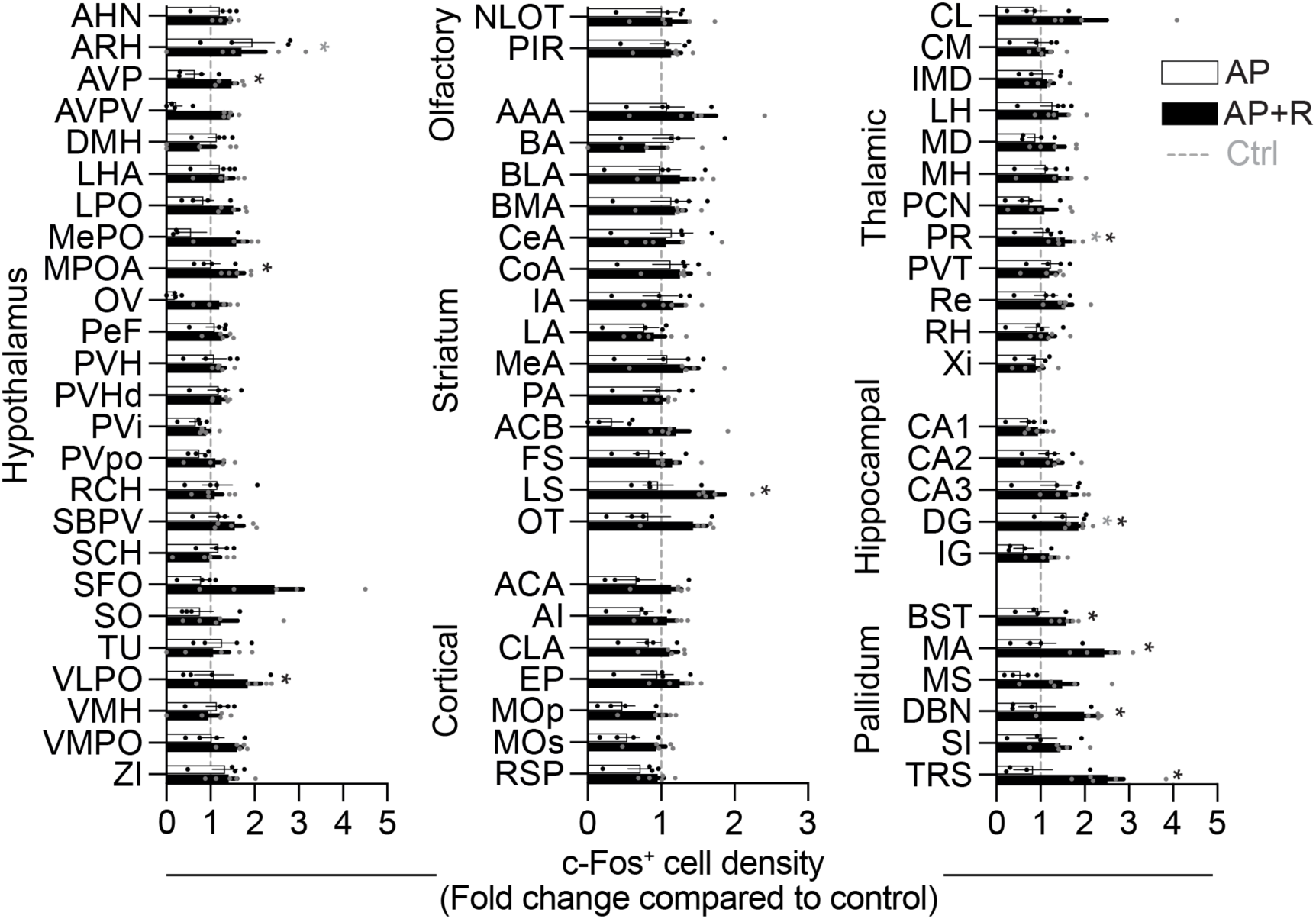
Recruitment of brain areas during juvenile alloparenting. c-Fos^+^ cell density in brain areas of P15 mice after alloparental behaviour (AP, n = 5 mice) and alloparental behaviour including pup retrieval (AP+R, n = 4 mice). Cell densities are normalised to non-exposed controls (Ctrl, n = 4 mice). Asterisks indicate significant differences between AP and controls (grey) or AP+R and controls (black). Statistics: *U* tests with *post hoc* Dunn’s test with Holm correction. Data are mean ± s.e.m.

**Extended Data Fig. 5:**
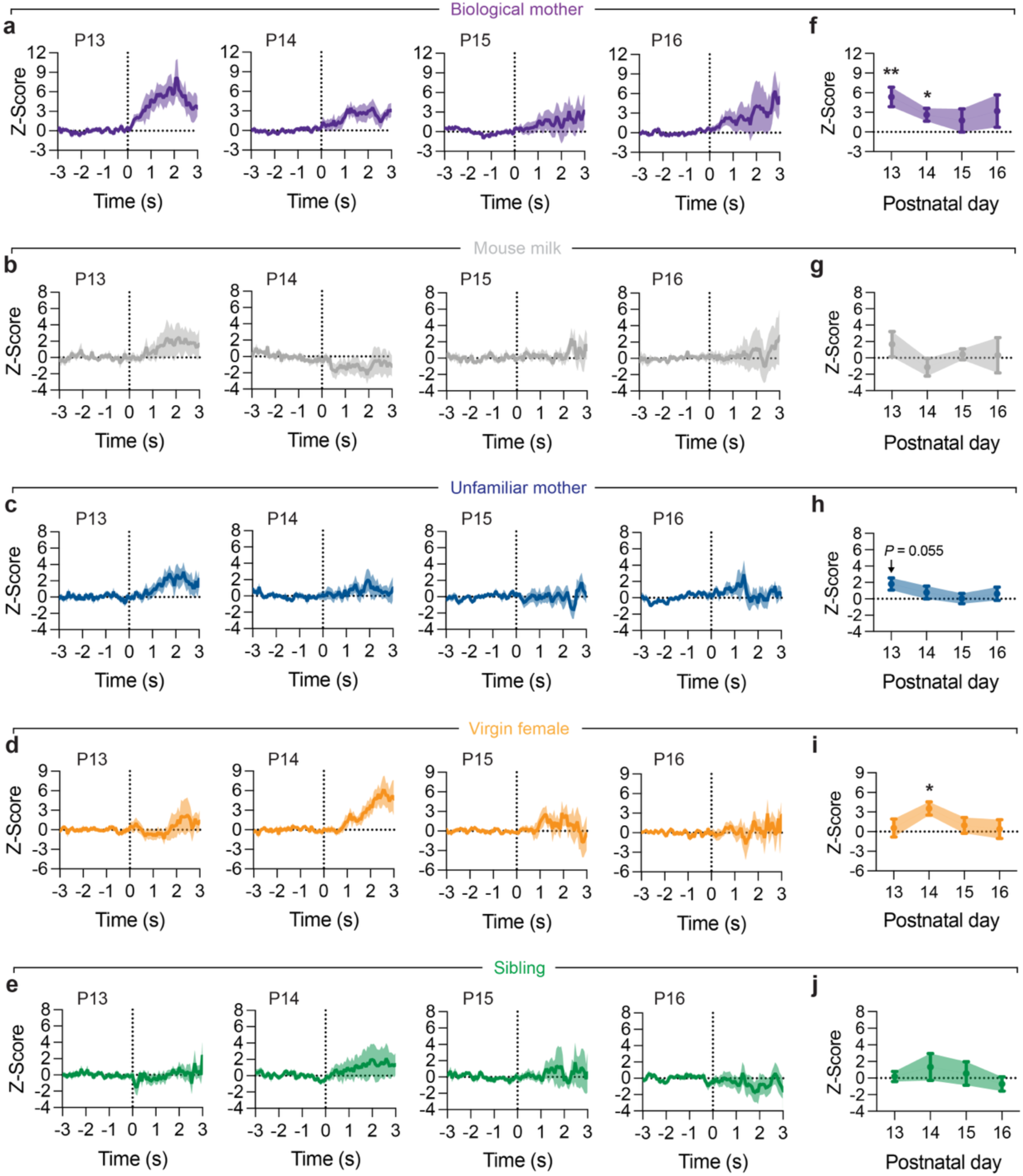
MPOA^Gal^ population activity during interactions with various stimuli. **a–j,** Peri-stimulus graphs showing MPOA^Gal^ neuron population response to interactions with biological mothers (**a**) and the averaged response vs baseline (**f**, see Methods, n = 9 mice), mouse milk (see Methods) (**b, g,** n = 6 mice), unfamiliar mothers (**c, h,** n = 10 mice), virgin females (**d, i,** n = 10 mice), and siblings (**e, j,** n = 10 mice). Dashed lines mark onset of interactions. Traces are mean ± SEM. Statistics: Wilcoxon signed-rank test with Benjamini-Hochberg FDR (**f–j**). Data are mean ± s.e.m. (error bars or shading).

**Extended Data Fig. 6:**
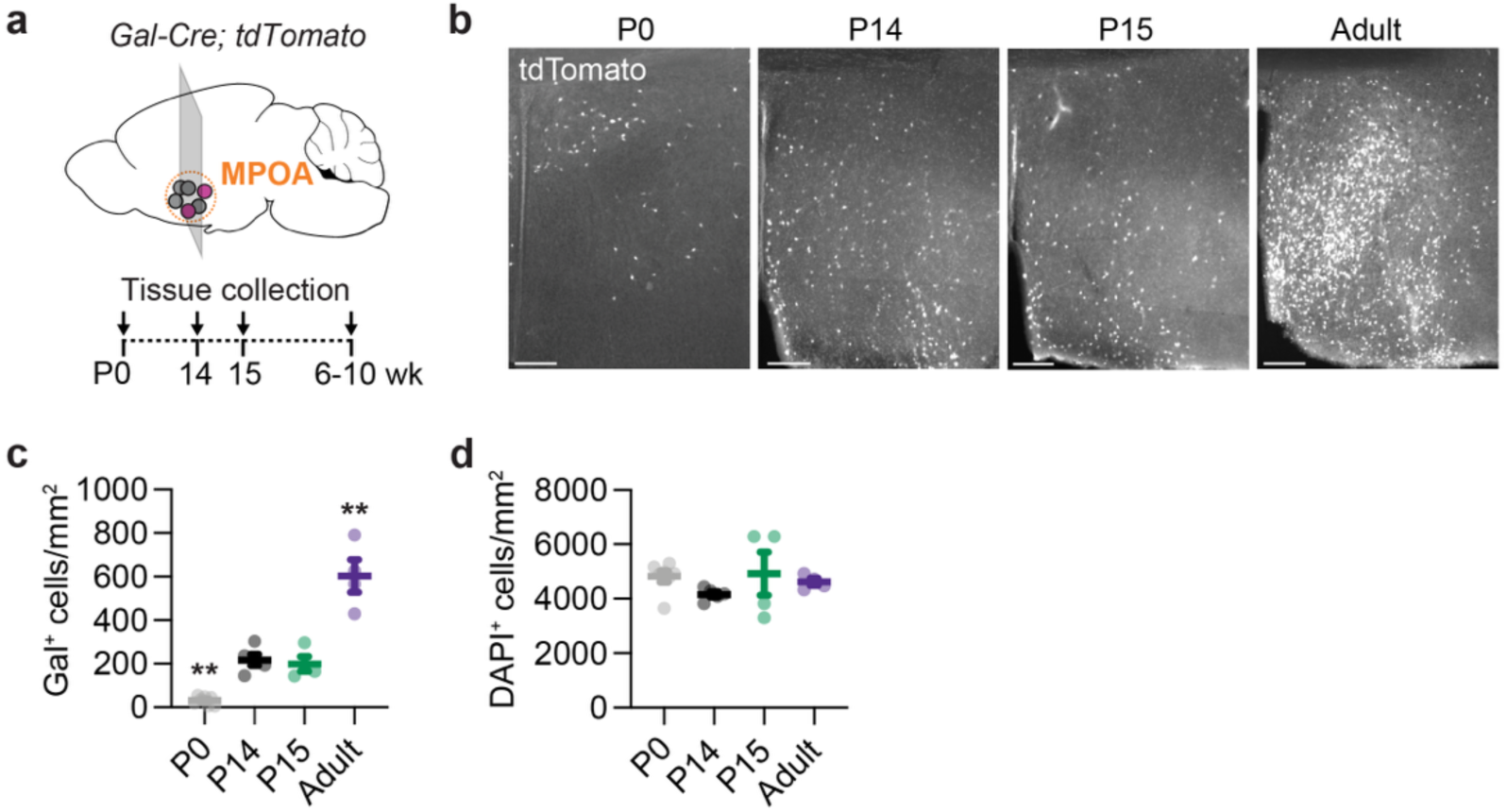
MPOA^Gal^ neuron density across postnatal development. **a,** Sampling time points to assess density of MPOA^Gal^ neurons in *Gal-Cre; tdTomato* mice. **b**, Example wide-field images of tdTomato-labelled MPOA^Gal^ neurons at P0, P14, P15 and in adulthood. Scale bars, 200 µm. **c**, Density of tdTomato-expressing MPOA^Gal^ neurons across age groups. Asterisks indicate significant changes compared to P14. **d,** Density of DAPI^+^ nuclei in MPOA across age groups (n = 7, 5, 4, 4 mice). Statistics: One-way ANOVA with *post hoc* Dunnett’s test (**c–d**). Data are mean ± s.e.m.

**Extended Data Fig. 7:**
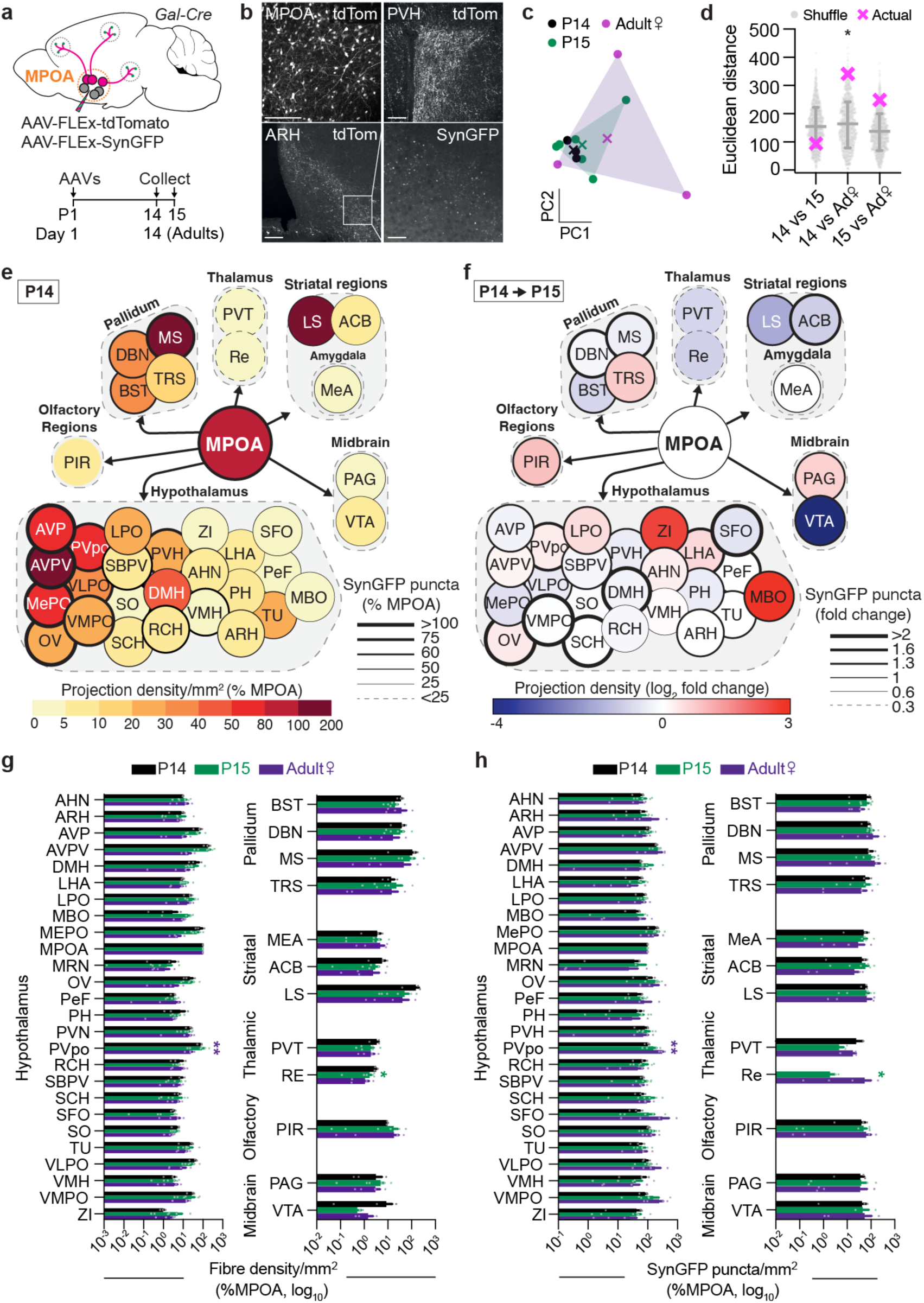
Brain-wide mapping of MPOA^Gal^ neuron projections. **a**, Anterograde tracing from MPOA^Gal^ neurons at P14, P15 and in adults. **b,** Example images of tdTomato-expressing MPOA^Gal^ neurons, tdTomato^+^ fibres in PVH and ARH, and SynGFP expression in ARH. Scale bar, 200 µm. **c,** Projection- and synapse distributions projected onto PC space. Each dot is an animal. **d,** Euclidean distance between MPOA^Gal^ input distributions at P14 (n = 5), P15 (n = 5) and 6–10-week-old virgin female adults (n = 4) in PC space, compared to shuffled (n = 1,000) distribution (mean ± SD). **e**, Average projection and synaptic density map at P14. **f,** Average fold change in projection and synaptic density between P14 and P15. **g**, Density of tdTomato-expressing fibres across brain areas. Data normalised to tdTomato expression in the MPOA. **h,** Density of SynGFP^+^ puncta across brain areas, normalised to SynGFP expression in the MPOA. Asterisks indicate significant changes between P14 and P15 (green) or adulthood (purple). Statistics: Permutation testing against shuffled data (**d**), unpaired *t*-test with *post hoc* Tukey’s HSD test for multiple comparisons (**g, h**), and *U* test with *post hoc* Dunn’s test with Holm correction (**g, h**). Data are mean ± s.e.m.

**Extended Data Fig. 8:**
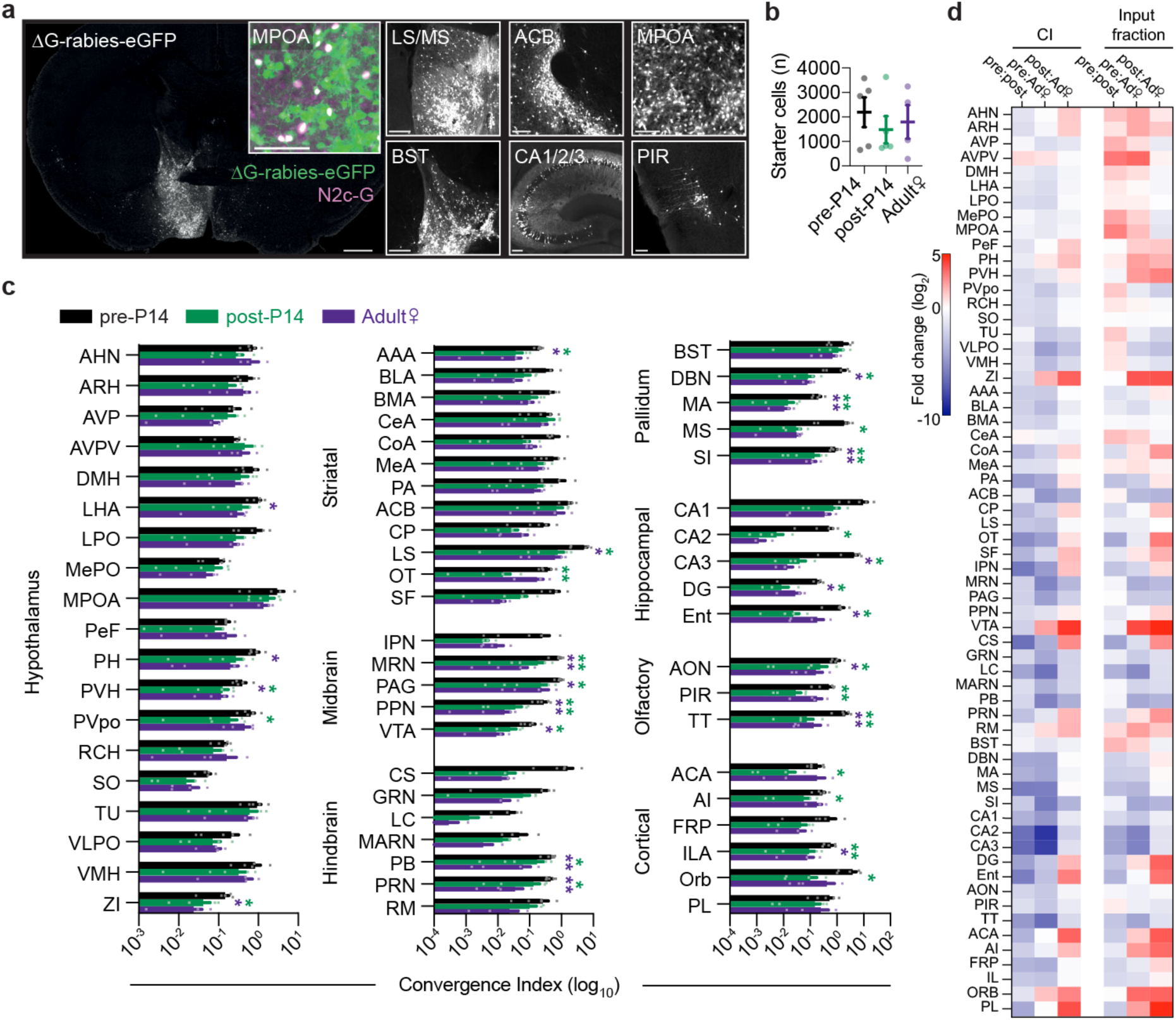
Brain-wide mapping of MPOA^Gal^ inputs in P14, P15 and adult mice. **a,** Example coronal section from a P14 mouse with rabies-positive neurons visible. Insets show starter cells as well as rabies-positive neurons in other brain regions. Scale bar, 1 mm, 200 µm (insets). **b,** Number of starter cells in MPOA in each group (n = 5, 5, 4 mice). **c,** Convergence index (ratio of number of rabies^+^ neurons per area to number of starter neurons in the MPOA, see Methods, n = 5, 5, 4 mice). Asterisks indicate significant differences between pre-P14 and post-P14 (green) or pre-P14 and adults (purple). **d,** Fold changes of convergence index and input fraction (see Methods) between pre-P14, post-P14 and adults (n = 5, 5, 4 mice). Statistics: One-way ANOVA with *post hoc* Dunnett’s test (**b**), unpaired *t*-test with *post hoc* Tukey’s HSD test for multiple comparisons (**c**), and *U* test with *post hoc* Dunn’s test with Holm correction (**c**). Data are mean ± s.e.m.

**Extended Data Fig. 9:**
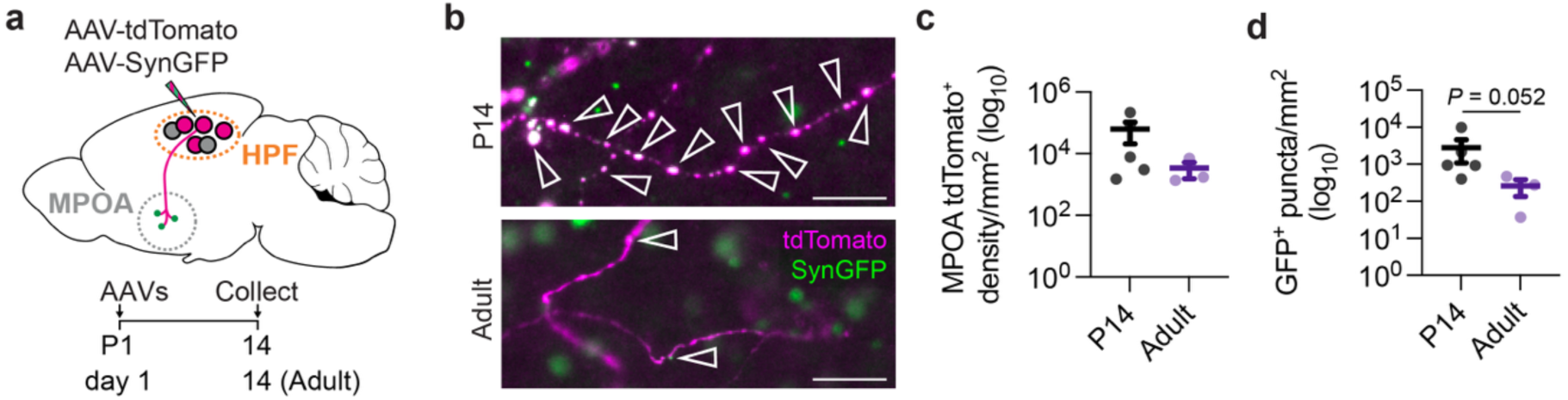
Higher synapse density of hippocampus→MPOA projections at P14 compared to adults. **a,** Anterograde tracing from the hippocampal formation (HPF) to the MPOA. **b,** Example images of tdTomato^+^ fibres in the MPOA of P14 and adults with SynGFP^+^, putative presynaptic contacts (arrowheads). Note that not all SynGFP^+^ puncta are highlighted in the P14 example. Scale bars, 10 µm. **c,** Density of tdTomato^+^ fibres in the MPOA. **d,** Density of SynGFP^+^ puncta in the MPOA (n = 5, 3 mice). Statistics: *U* test with *post hoc* Dunn’s test with Holm correction (**c, d**). Data are mean ± s.e.m.

**Extended Data Fig. 10:**
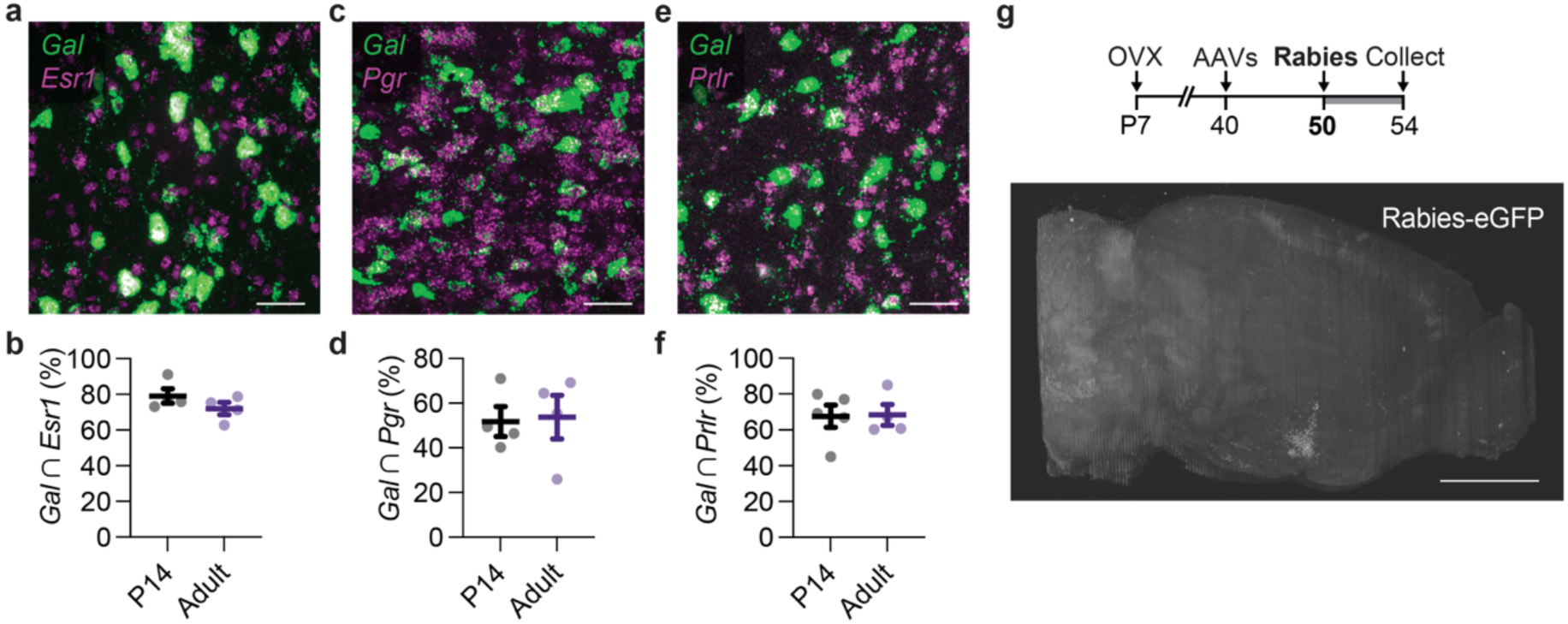
MPOA^Gal^ hormone receptors are comparable between juvenile and adult. **a–f:** MPOA^Gal^ receptors for estrogen (*Esr1,* **a–b**), progesterone (*Pgr,* **c–d**) and prolactin (*Prlr,* **e–f**) are the same between P14 and adult female virgin mice. smFISH examples (**a, c, e**) shown above quantifications (**b, d, f**). **g,** Timeline of surgical procedures (top) and side view of serial section two-photon tomography images (bottom) with rabies-positive neurons in a juvenile ovariectomy (OVX) animal. Statistics: *U* test with *post hoc* Dunn’s test with Holm correction (**b, d, f**). Data are mean ± s.e.m.

**Extended Data Fig. 11:**
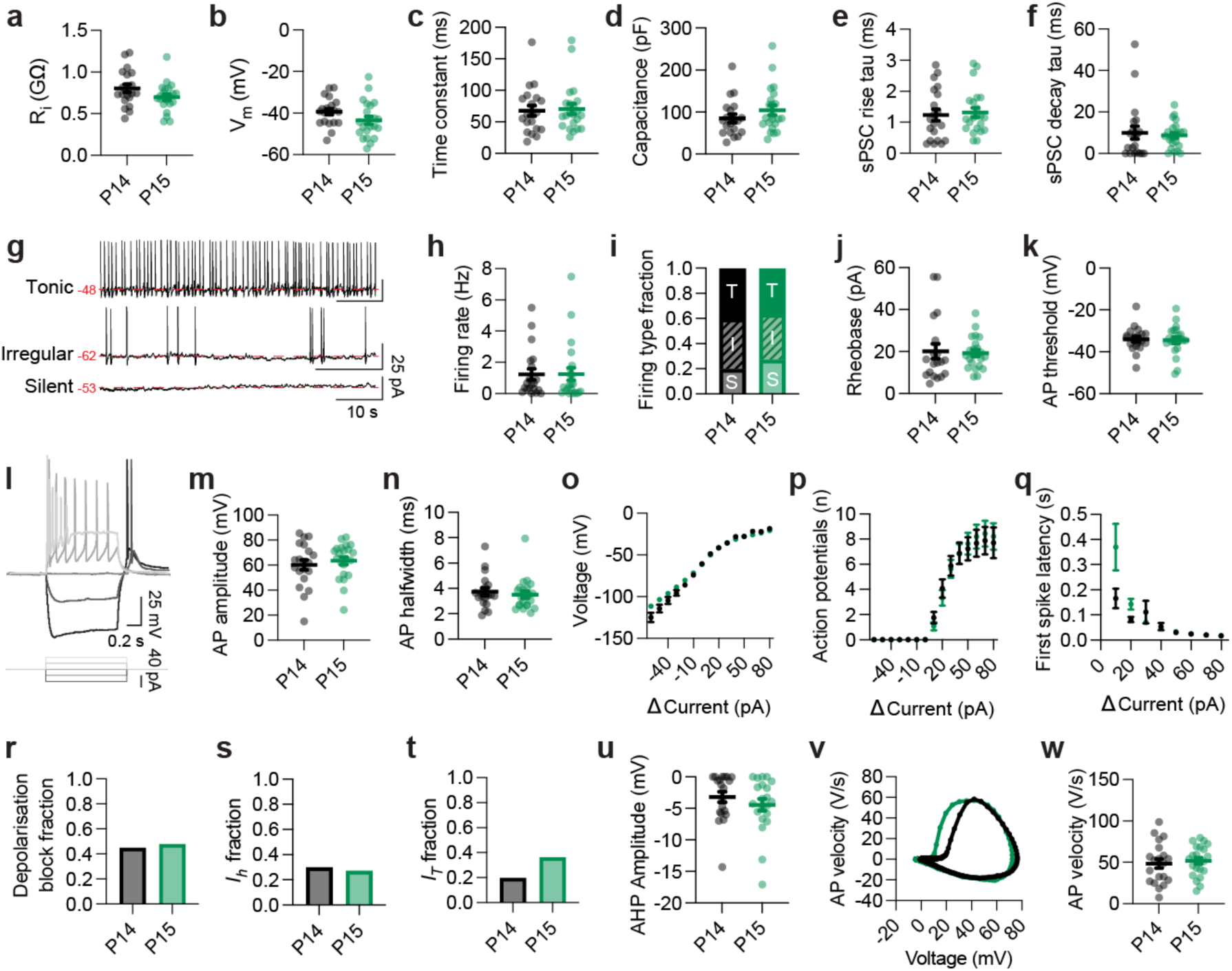
Biophysical properties of MPOA^Gal^ neurons at P14 and P15. Characterisation of MPOA^Gal^ electrophysiological properties at P14 and P15 (20, 22, neurons from n = 5, 5 mice) (**a–f**): Input resistance (**a**), resting membrane potential (**b**), time constant (**c**), capacitance (**d**), spontaneous postsynaptic current (sPSC) rise time constant (**e**), sPSC decay time constant (**f**). **g**, Example recording traces of MPOA^Gal^ baseline activity patterns. Red dashed line indicates baseline voltage (mV). **h,** Action potential (AP) firing rate at baseline. **i,** Spontaneous activity types of MPOA^Gal^ neurons at baseline. T, tonic, I, irregular, S, silent. **j**, Rheobase input current required to elicit an action potential from baseline. **k**, membrane potential at which the first AP is fired. **l,** Example trace of a neuron receiving current steps from –40 to 40 pA at 20 pA intervals (top). Current injections corresponding to voltage trace. Colours correspond to the voltage response above (bottom). **m–w,** Electrophysiological properties corresponding to the voltage response to 20 pA current injections: Amplitude of the first AP fired in response to a 20 pA current step (**m**), halfwidth of the first AP fired in response to a 20 pA current step (**n**), membrane voltage in response to current steps (**o**), APs elicited by current steps (**p**), first spike latency of APs elicited by current steps (**q**), proportion of neurons going into depolarisation block with sequential current steps (**r**), proportion of neurons with *Ih* current in response to a hyperpolarising –60 pA current step (**s**), proportion of neurons with *IT* current in response to a hyperpolarising –60 pA current step (**t**), after-hyperpolarising potential (AHP) amplitude of the first action potential (AP) fired in response to a 20 pA current step (**u**), phase plot of the AP velocity per voltage of the first AP fired in response to a 20 pA current step (**v**), maximal AP velocity of the first AP fired in response to a 20 pA current step (**w**). Statistics: *U* tests with *post hoc* Dunn’s test with Holm correction (**a–f, h, j–k, m–q, u, w**); Chi-square test (**i**), Fisher’s exact test with *post hoc* Benjamini-Hochberg FDR correction (**r–t**). Data are mean ± s.e.m.

**Extended Data Fig. 12:**
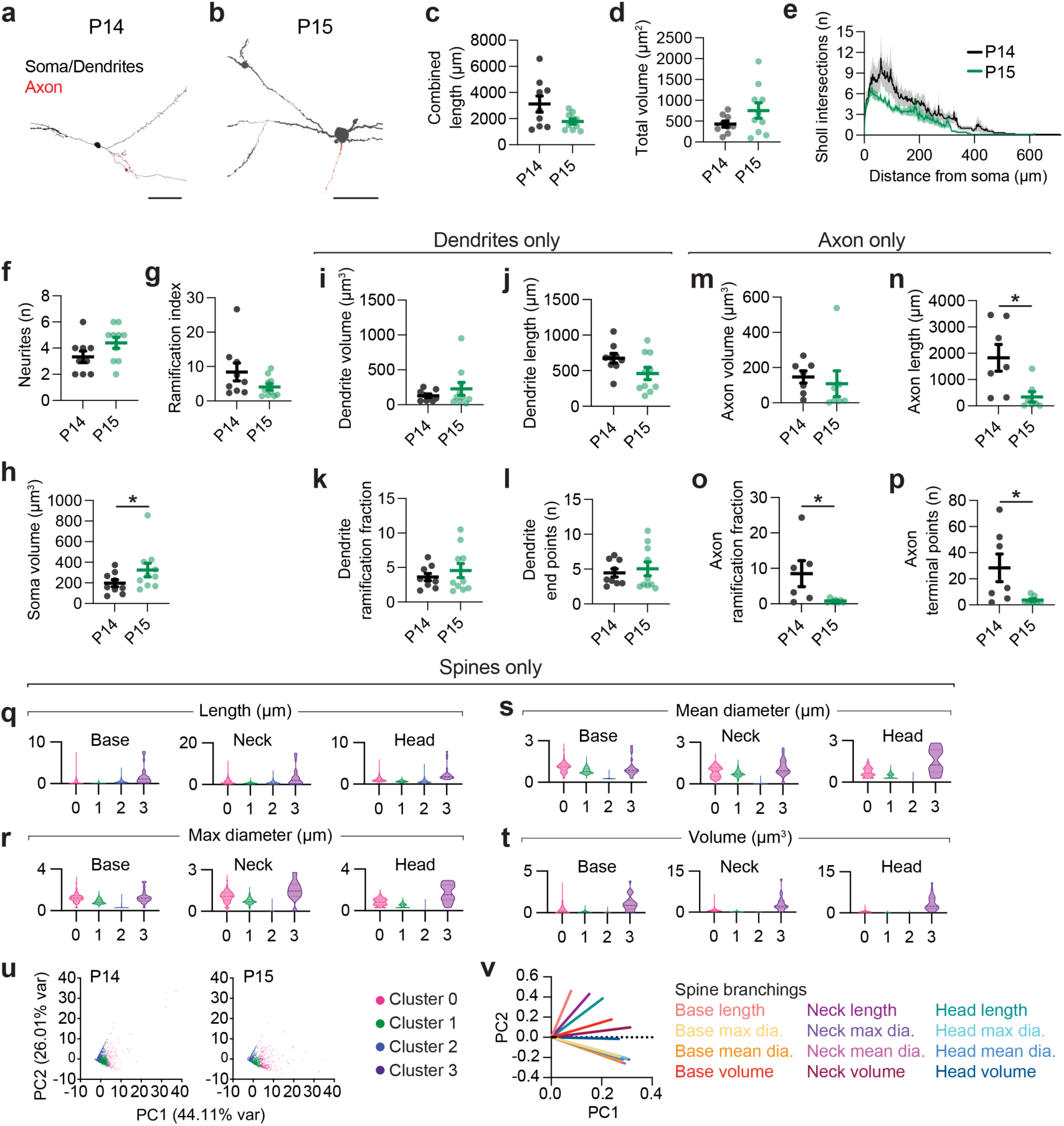
Morphological properties of MPOA^Gal^ neurons at P14 and P15. **a–b**, Example MPOA^Gal^ neuron reconstructions from P14 (**a**) and P15 (**b**) mice. Scale bars, 50 µm. **c–h,** Total projection length (**c**), neuronal volume (**d**), Sholl analysis (**e**), neurites (**f**), ramification index (**g**), and soma volume (**h**) (9, 10 neurons from n = 6, 5 mice; for definitions see Methods). **i– p,** dendritic (**i–l**; 9, 10 neurons from 6, 5 mice) and axonal (**m–p**; 7, 7 neurons from 4, 5 mice) morphological parameters at P14 and P15 (average values per neuron are shown): volume (**i, m**), length (**j, n**), ramification fraction (**k, o**), end points (**l, p**). **q–t**, Parameter metrics involved in PCA for each spine cluster divided by lower (base) middle (neck) and upper (head) third of spine length: length (**q**), maximum diameter (**r**), volume (**s**), mean diameter (**t**). **u**, K-means clustering of dendritic spines from P14 and P15 projected in PC space (2,488, 1,549, spines from 9, 10 neurons, n = 6, 5). Example spines from each cluster showing in Fig. 2q. **v**, Radar plot showing relative contribution of each spine parameter to PC1 and PC2. Statistics: *U* tests with *post hoc* Dunn’s test with Holm correction (**c–p**). Data are mean ± s.e.m. (error bars or shading).

**Extended Data Fig. 13:**
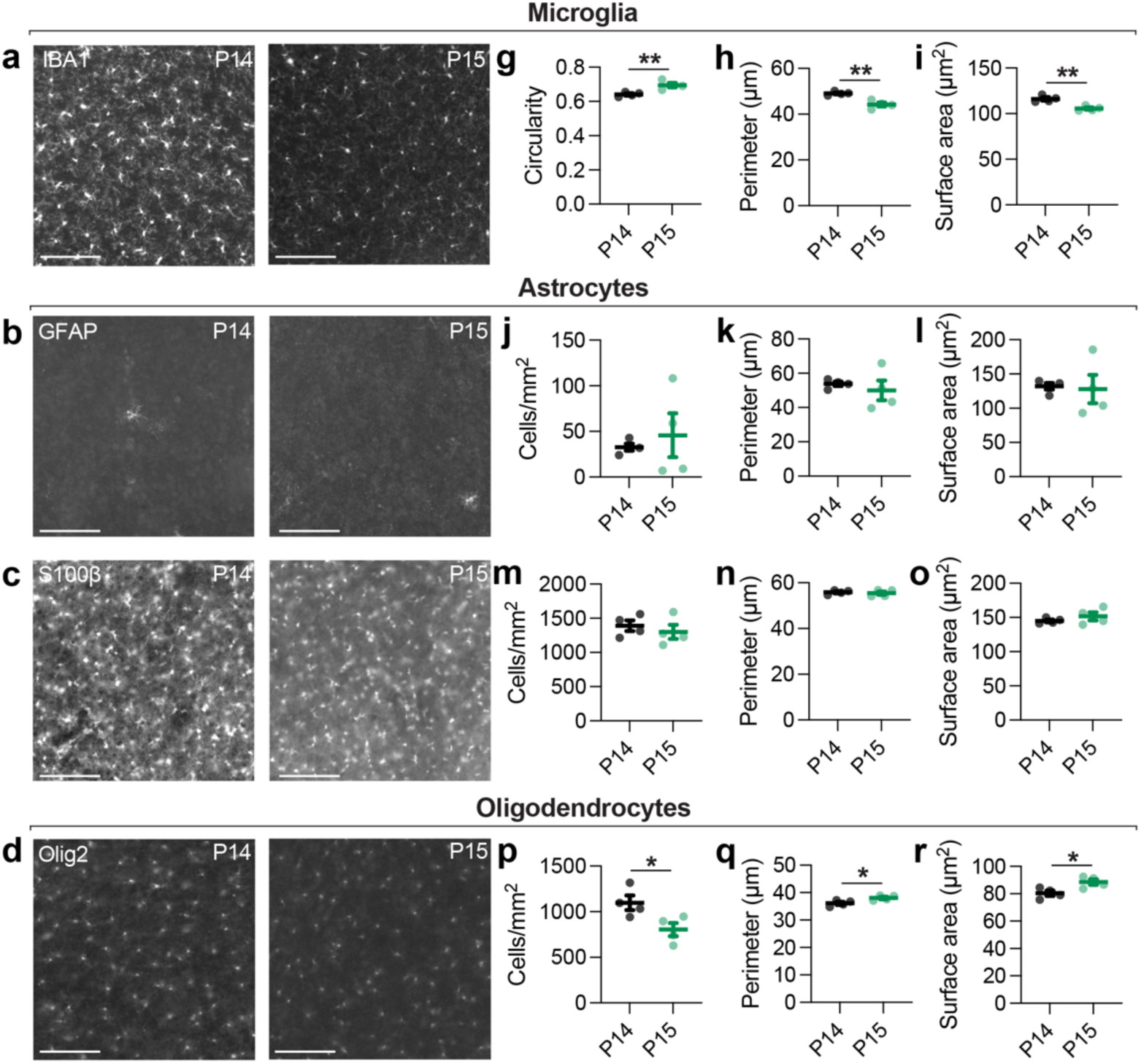
Density and morphological properties of MPOA glial cells between P14–15. **a–d,** Wide-field images of immunostained MPOA glia cell types: microglia (**a**), astrocytes (**b– c**) and oligodendrocytes (**d**). Scale bars, 100 µm. **g–r,** Morphological parameters of immunostained microglia (**g–i,** n = 4, 4 mice), astrocytes (**j–o,** n = 4, 4 mice) and oligodendrocytes (**p–r,** n = 4, 4 mice). Statistics: *U* tests with *post hoc* Dunn’s test with Holm correction (**g–r**). Data are mean ± s.e.m.

**Extended Data Fig. 14:**
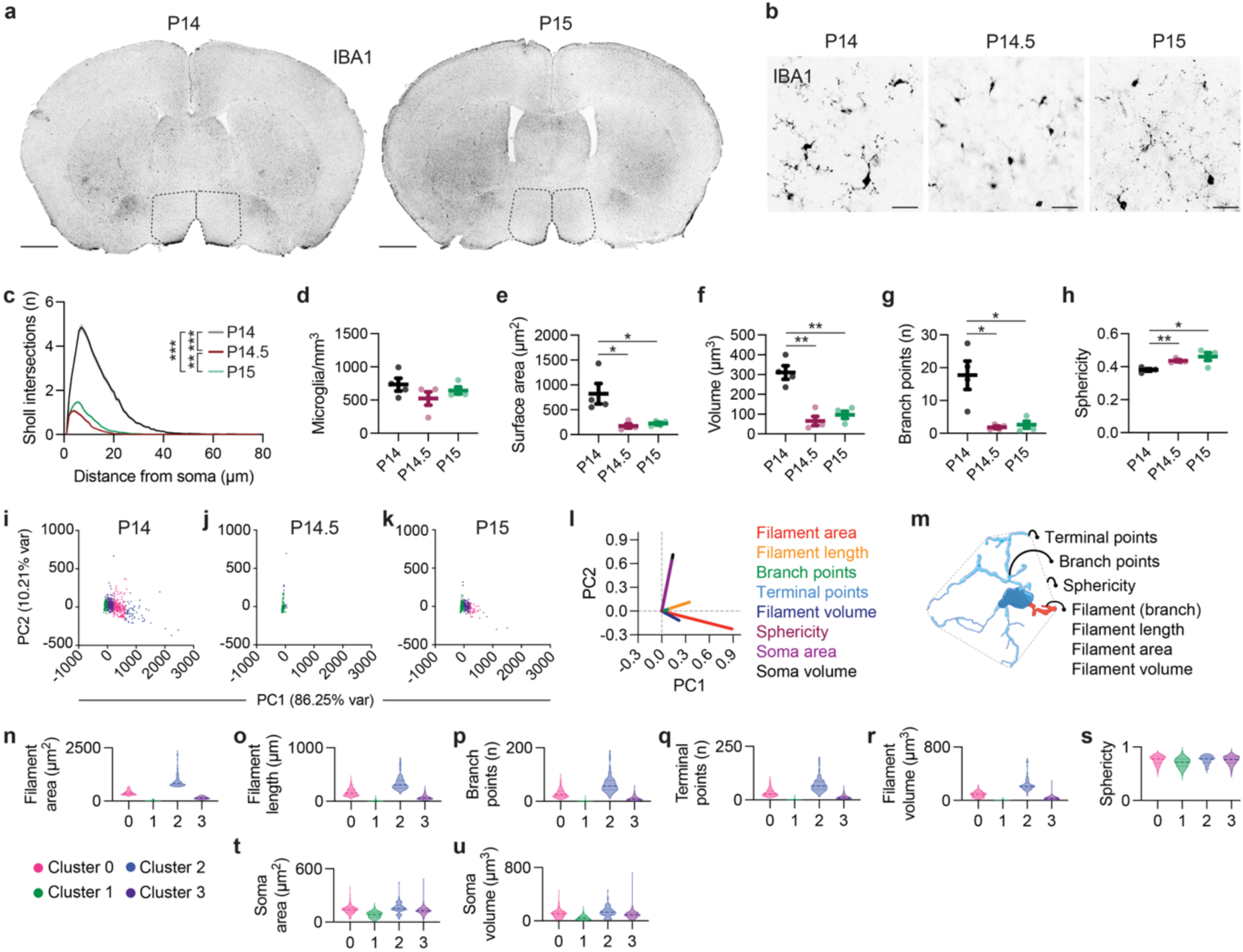
MPOA microglia have altered morphology at P14.5. **a,** Wide-field brain slice images of IBA1-stained microglia at P14 and P15. MPOA outlined. Scale bars, 1 mm. **b,** Confocal images of IBA1-stained MPOA microglia at P14, P14.5 and P15. Scale bars, 100 µm. **c–h,** Morphological parameters of MPOA microglia across age groups (averages from n = 4, 4, 4 mice): Sholl analysis (**c**), MPOA microglia density (**d**), microglia surface area (**e**), volume (**f**), branch points (**g**, see panel **m**), and sphericity (**h**, see panel **m**). **i–k**, K-means clustering of MPOA microglial morphologies at P14 (**i**), P14.5 (**j**) and P15 (**k**) projected in PC space (586, 359, 607 microglia from n = 4, 4, 4 mice, see Methods). Example microglia from each cluster showing in Fig. 3f. **l**, Radar plot showing the relative contribution of each morphological parameter to the first two PCs. **m,** Example reconstructed microglia (Cluster 2) with morphological parameters highlighted. **n–u**, Distribution of morphological parameters across microglia clusters. Statistics: One-way ANOVA with Holm correction (**c–h**). Data are mean ± s.e.m.

**Extended Data Fig. 15:**
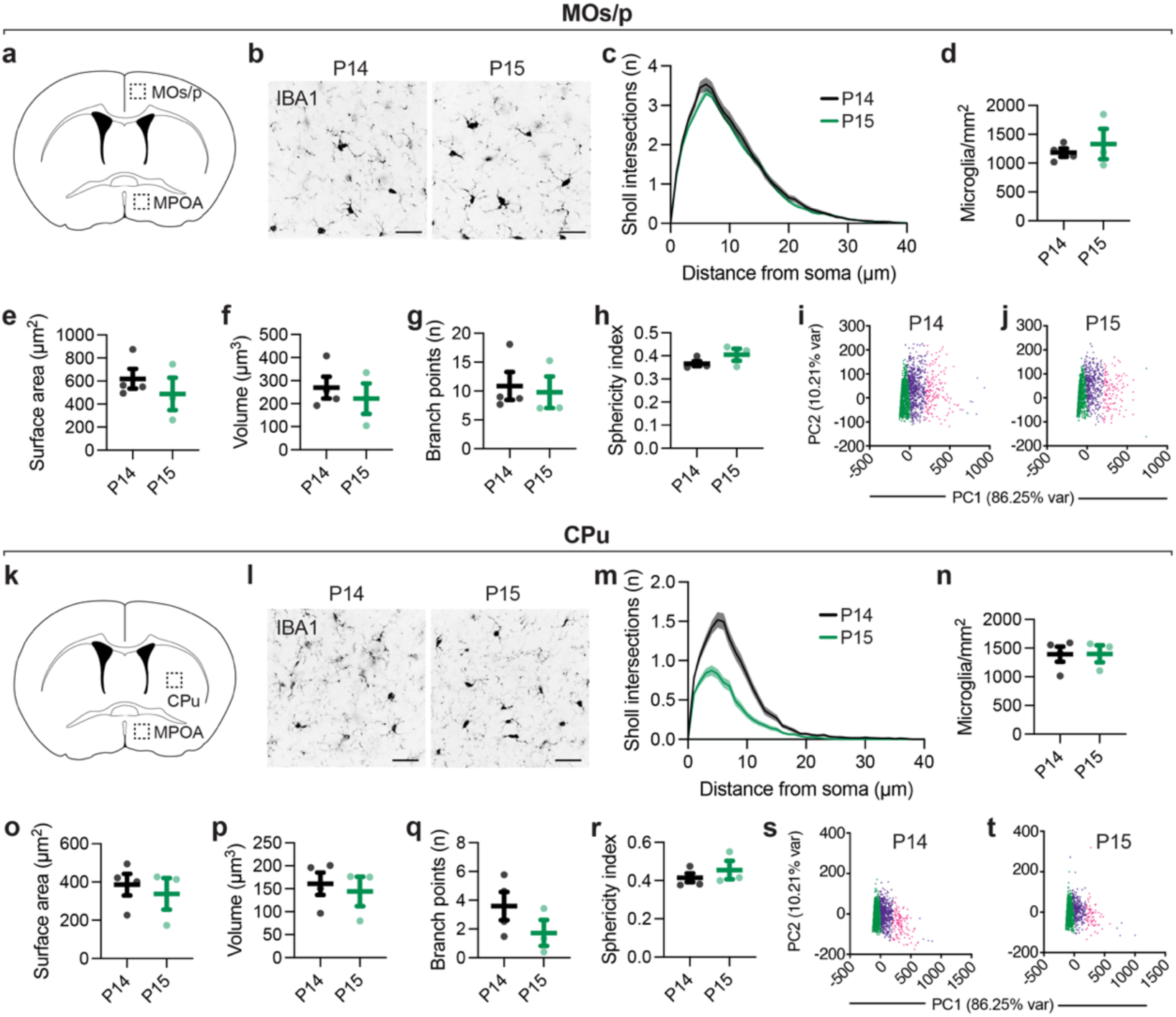
Microglia in motor cortex and striatum do not undergo morphological changes between P14–15. **a, k,** Cartoon brain slice showing the location of MOs/p (**a**) and CPu (**k**) relative to the MPOA. **b, l,** Confocal images of IBA1-stained microglia at P14 and P15 in MOs/p (**b**) and CPu (**l**). Scale bars, 50 µm. **c–h, m–r,,** Morphological parameters of microglia in MOs/p (**c–h;** averages from n = 4, 3 mice) and CPu (**m–r;** averages from n = 4, 3 mice) at P14 and P15: Sholl intersections (**c, m**), microglia density (**d, n**), surface area (**e, o**), volume (**f, p**), number of branch points (**g, q**), and sphericity (**h, r**). **i–j, s–t,** K-means clustering of microglial morphologies in MOs/p (**i–j,** 1,844, 1,220 microglia from n = 4, 3 mice) and CPu (**s–t,** 2,274, 1,787 microglia from n = 4, 3 mice). Example microglia from each cluster showing in Fig. 3f. Statistics: *U* tests with *post hoc* Dunn’s test with Holm correction (**c–h, m–r**). Data are mean ± s.e.m. (error bars or shading).

**Extended Data Fig. 16:**
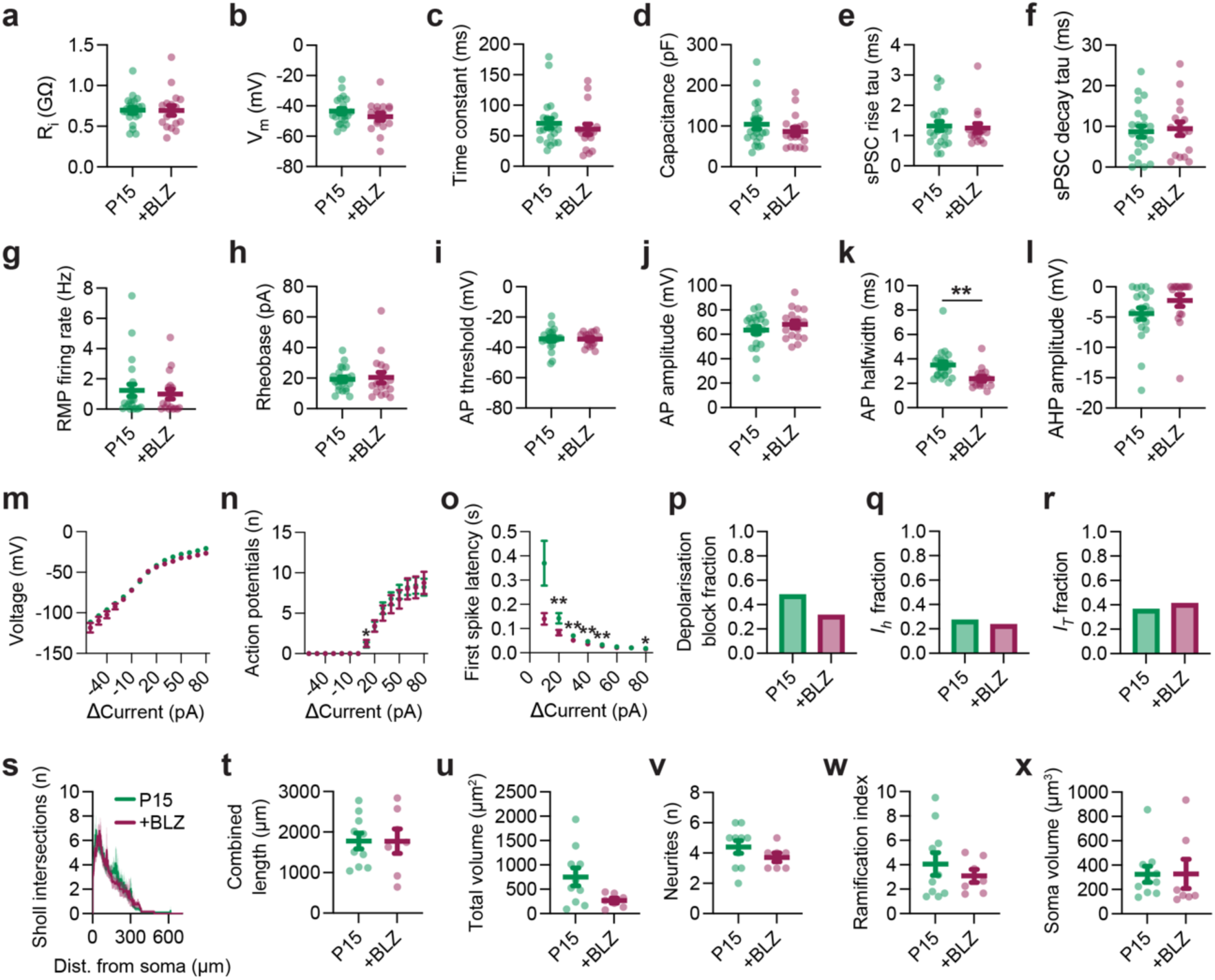
Biophysical and morphological properties of MPOA^Gal^ neurons from control and BLZ-treated P15 mice. **a–r,** Biophysical parameters of MPOA^Gal^ neurons from controls and BLZ-treated mice (22, 18 neurons from n = 5, 4 mice): resting membrane potential (**a**), input resistance (**b**), time constant (**c**), capacitance (**d**), sPSC rise time constant (**e**), sPSC decay time constant (**f**), baseline firing rate (**g**), rheobase input current required to elicit a AP from baseline (**h**), membrane potential at which the first AP is fired (**i**), amplitude of the first AP fired in response to a 20 pA current step (**j**), halfwidth of the first AP fired in response to a 20 pA current step (**k**), after-hyperpolarising potential (AHP) amplitude of the first AP fired in response to a 20 pA current step (**l**), membrane voltage in response to current steps (**m**), APs elicited by current steps (**n**), first spike latency of APs elicited by current steps (**o**), proportion of neurons going into depolarisation block with sequential current steps (**p**), proportion of neurons with *Ih* current in response to a hyperpolarising –60 pA current step (**q**), proportion of neurons with *IT* current in response to a hyperpolarising –60 pA current step (**r**). **s–x**, Biophysical parameters of MPOA^Gal^ neurons from controls and BLZ-treated mice (10, 7 neurons from n = 5, 5 mice): Sholl analysis (**s**), total projection length (**t**), neuronal volume (**u**), neurites (**v**), ramification index (**w**), and soma volume (**x**). Statistics: *U* tests with *post hoc* Dunn’s test with Holm correction (**a–o, s–x**), Fisher’s exact test with *post hoc* Benjamini-Hochberg FDR test (**p–r**). Data are mean ± s.e.m. (error bars or shading).

**Extended Data Fig. 17:**
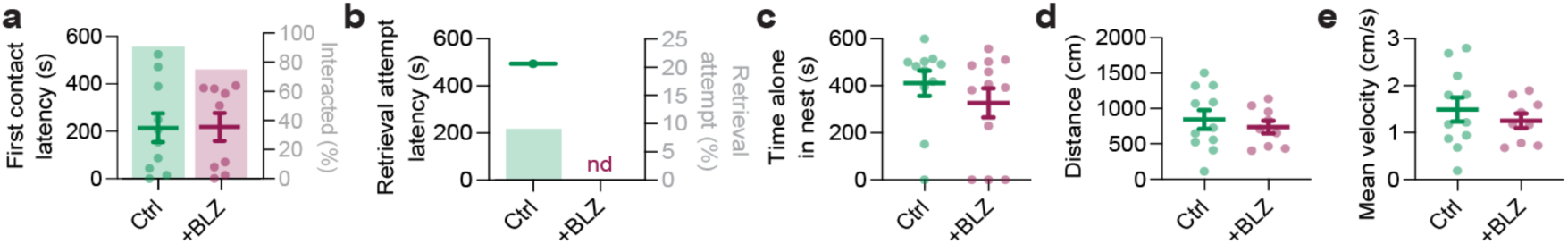
Characterisation of juvenile alloparental behaviour in microglia-depleted mice. Pup interactions (**a–c,** n = 11, 12 mice) and locomotion (**d–e,** n = 11, 9 mice) in controls and BLZ-treated mice at P15: pup contact latency (left) and percentage of mice interacting with pups (right) (**a**), retrieval attempt latency (left) and percentage of mice attempting retrievals (right, **b**), time spent in nest (**c**), distance moved (**d**) and mean velocity in 10-min open field assay (**e**). Statistics: Fisher’s exact test with *post hoc* Benjamini-Hochberg FDR correction (**a, b**), *U* tests with *post hoc* Dunn’s test with Holm correction (**a, c–e**), Kaplan-Meier survival analysis log-rank test (**b**). Data are mean ± s.e.m.

**Extended Data Table 1:**
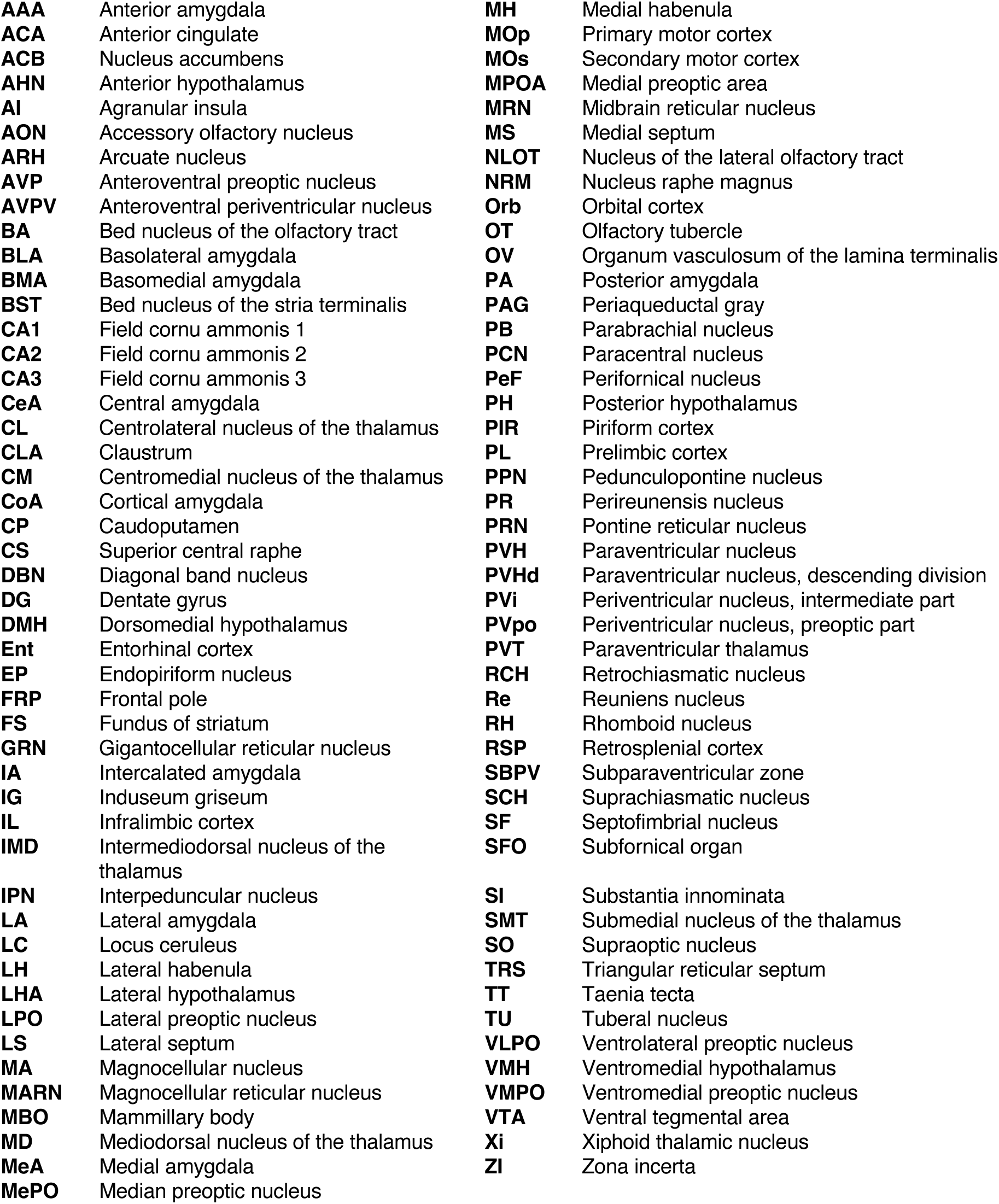
Abbreviations of brain regions.

|  |  |  |  |
| --- | --- | --- | --- |
| **AAA** | Anterior amygdala | **MH** | Medial habenula |
| **ACA** | Anterior cingulate | **MOp** | Primary motor cortex |
| **ACB** | Nucleus accumbens | **MOs** | Secondary motor cortex |
| **AHN** | Anterior hypothalamus | **MPOA** | Medial preoptic area |
| **AI** | Agranular insula | **MRN** | Midbrain reticular nucleus |
| **AON** | Accessory olfactory nucleus | **MS** | Medial septum |
| **ARH** | Arcuate nucleus | **NLOT** | Nucleus of the lateral olfactory tract |
| **AVP** | Anteroventral preoptic nucleus | **NRM** | Nucleus raphe magnus |
| **AVPV** | Anteroventral periventricular nucleus | **Orb** | Orbital cortex |
| **BA** | Bed nucleus of the olfactory tract | **OT** | Olfactory tubercle |
| **BLA** | Basolateral amygdala | **OV** | Organum vasculosum of the lamina terminalis |
| **BMA** | Basomedial amygdala | **PA** | Posterior amygdala |
| **BST** | Bed nucleus of the stria terminalis | **PAG** | Periaqueductal gray |
| **CA1** | Field cornu ammonis 1 | **PB** | Parabrachial nucleus |
| **CA2** | Field cornu ammonis 2 | **PCN** | Paracentral nucleus |
| **CA3** | Field cornu ammonis 3 | **PeF** | Perifornical nucleus |
| **CeA** | Central amygdala | **PH** | Posterior hypothalamus |
| **CL** | Centrolateral nucleus of the thalamus | **PIR** | Piriform cortex |
| **CLA** | Clastrum | **PL** | Prelimbic cortex |
| **CM** | Centromedial nucleus of the thalamus | **PPN** | Pedunculo pontine nucleus |
| **CoA** | Cortical amygdala | **PR** | Perireunensis nucleus |
| **CP** | Caudoputamen | **PRN** | Pontine reticular nucleus |
| **CS** | Superior central raphe | **PVH** | Paraventricular nucleus |
| **DBN** | Diagonal band nucleus | **PVHd** | Paraventricular nucleus, descending division |
| **DG** | Dentate gyrus | **PVi** | Periventricular nucleus, intermediate part |
| **DMH** | Dorsomedial hypothalamus | **PVpo** | Periventricular nucleus, preoptic part |
| **Ent** | Entorhinal cortex | **PVT** | Paraventricular thalamus |
| **EP** | Endopiriform nucleus | **RCH** | Retrochiasmatic nucleus |
| **FRP** | Frontal pole | **Re** | Reuniens nucleus |
| **FS** | Fundus of striatum | **RH** | Rhomboid nucleus |
| **GRN** | Gigantocellular reticular nucleus | **RSP** | Retrosplenial cortex |
| **IA** | Intercalated amygdala | **SBPV** | Subparaventricular zone |
| **IG** | Induseum griseum | **SCH** | Suprachiasmatic nucleus |
| **IL** | Infralimbic cortex | **SF** | Septofimbrial nucleus |
| **IMD** | Intermediodorsal nucleus of the thalamus | **SFO** | Subfornical organ |
| **IPN** | Interpeduncular nucleus | **SI** | Substantia innominata |
| **LA** | Lateral amygdala | **SMT** | Submedial nucleus of the thalamus |
| **LC** | Locus ceruleus | **SO** | Supraoptic nucleus |
| **LH** | Lateral habenula | **TRS** | Triangular reticular septum |
| **LHA** | Lateral hypothalamus | **TT** | Taenia tecta |
| **LPO** | Lateral preoptic nucleus | **TU** | Tuberal nucleus |
| **LS** | Lateral septum | **VLPO** | Ventrolateral preoptic nucleus |
| **MA** | Magnocellular nucleus | **VMH** | Ventromedial hypothalamus |
| **MARN** | Magnocellular reticular nucleus | **VMPO** | Ventromedial preoptic nucleus |
| **MBO** | Mammillary body | **VTA** | Ventral tegmental area |
| **MD** | Mediodorsal nucleus of the thalamus | **Xi** | Xiphoid thalamic nucleus |
| **MeA** | Medial amygdala | **ZI** | Zona incerta |
| **MePO** | Median preoptic nucleus |  |  |

**Supplementary Video 1:** Example of juvenile alloparental behaviour. A P15 male retrieves pups to the nest and engages in pup grooming.

## Methods

### Ethical Approval

All animal procedures performed in this study were approved by the United Kingdom Government (Home Office) and by the Crick Institutional Animal Welfare Ethical Review Panel.

### Animals

Mice were housed in individually ventilated cages on a 12 h: 12 h light:dark cycle (light from 22:00–10:00 h) with *ad libitum* access to food and water. The *Gal-Cre* BAC transgenic line (Stock: Tg(*Gal::cre*) KI87Gsat/Mmucd, 031060-UCD) was imported from the Mutant Mouse Regional Resource Center. This line was crossed to Cre-dependent Rosa26 tdTomato (Ai9, JAX #007909) reporter mice to visualise Gal-expressing neurons. Sex was determined at birth by the presence of pigmentation on the genital ridge in males, which was absent in females^63^.

### Pup-directed behaviour assay

Experiments were performed in the first half of the dark phase. Juvenile mice were tested from P13 at the earliest, as this is when eye opening approximately begins^24^. For the testing cage, home cage bedding was spread onto new cage bedding. Home cage nesting material was placed in the corner of the test cage. Mice were transferred to the test cage and left for 10–15 min to habituate. To compare alloparental behaviour between juveniles and adults, pup interactions of female mice were assessed at P15 and again at P80. Adult mice were group-housed in same-sex cages of 3–5 mice. Males were group-housed until behavioural testing to promote spontaneous parental behaviour. Females were individually housed for 4 d before behavioural testing as in previous work^64^. Two P0-2 foster pups from C57BL/6J litters were randomly chosen and placed in corners opposite to the nesting. Interactions between test mouse and pups were recorded for 10 min with a camera (Basler Ace GigE, acA1300-60gmNIR) capturing a top view of the testing cage at a 30 Hz framerate. Manual scoring of the behaviour was carried out using EthoVision XT 14 (Noldus). Duration of behaviour was calculated as the total behavioural period over 600 s. Time alone in nest was defined as the mouse being alone within the boundaries of the nesting material. Contact latency was defined as the time from introduction of the first pup to the test cage until the first contact of the pup with the test mouse’s nose. Alloparental interactions were defined as close contact with a pup involving licking, pup displacement and rhythmic head movements. Retrieval attempt latency was defined as the time from test onset to when the first pup was picked up in the mouth and carried in the direction of the nest. Interacted (%) or Retrieval attempt (%) refers to the percentage of mice that either interacted with the pup or attempted to retrieve or successfully retrieved the pup to the nest, respectively. For chemogenetic experiments, mice were injected with 0.1 mg/kg DCZ (HelloBio, HB9126) prior to habituation.

#### c-Fos experiments

For c-Fos experiments, P15 mice were scored as alloparental (AP) or alloparental and retrieving (AP+R) based on whether pup retrieval (or a retrieval attempt) occurred. Ninety min after the first interaction with a pup, mice were transcardially perfused with 20 mL phosphate-buffered saline (PBS) followed by 20 mL paraformaldehyde (PFA, 4% in PBS). Control mice were left in the test cage for 10 min without introducing pups and perfused 90 min later. Brains were then dissected out and post-fixed overnight in 4% PFA before being washed in PBS. c-Fos immunostaining was carried out as described below. For smFISH experiments, P15 mice were transcardially perfused with 20 mL phosphate-buffered saline (PBS) 45 min after the first interaction with a pup, and brains dissected and embedded in OCT (Tissue-Tek, 4583), frozen on dry ice and stored at –80°C. smFISH was carried out as described below.

#### Novel stimulus preference testing

For place preference testing, a novel cage with clean bedding was used as the test arena. One pup (P0–2) and one toy Lego brick (32 × 16 mm) were placed at opposite ends of the cage. A juvenile that had not previously been exposed to younger pups was placed in the middle of the cage equidistant from the pup and brick. Interactions between the test mouse and stimuli were recorded for 10 min with a camera (Basler Ace GigE, acA1300-60gmNIR) capturing a top view of the testing cage at a 30 Hz frame rate. Manual scoring of the time the test mouse was in contact with either the pup (t_p_) or the brick (t_b_) was carried out in EthoVision XT 14 (Noldus). Preference was then determined as follows:

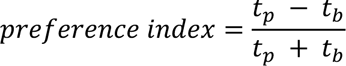

### Juvenile developmental scoring

Developmental scoring was based on parameters set out by Hill *et al*., (2008)^24^. Briefly, P14 and P15 mice were weighed and checked for whether their eyes were open. Auditory startle responses were assessed using a loud clap (∼90 dB), 10 cm from the subject’s location. Mice were held up to grasp a metal rod, left suspended 7 cm over nesting material and the time from grasp to falling was recorded. The forelimb grasp test lasted for 10 s. For locomotory assessment, the distance and speed of mice in an empty cage with standard bedding was recorded for 10 min and quantified using automatic tracking in EthoVision XT 14 (Noldus).

### Histology and immunostaining

Mice were taken during the first half of the dark phase and transcardially perfused with 20 mL phosphate-buffered saline (PBS) followed by 20 mL paraformaldehyde (PFA, 4% in PBS). Brains were then dissected out and post-fixed overnight in 4% PFA before being washed in PBS. 60 µm coronal sections were cut using a vibratome (Leica VT1000S) and either immediately mounted to Superfrost Plus slides (VWR, 48311-703) with DAPI-containing Vectashield mounting medium (Vector Laboratories, H-1200); or kept for immunostaining. For immunostaining, slices were kept in individual wells in 48-well culture plates and washed in blocking buffer (0.3% Triton X-100, 3% BSA, 2% normal goat serum in PBS). Incubation with primary antibody was performed in blocking buffer, for 48 h on a nutator at 4°C, with the following antibodies: chicken-anti-GFP (Abcam AB13970; 1:1,000), rabbit-anti-RFP (Rockland Immunochemicals, 600-401-379; 1:1,000); rabbit-anti-IBA1 (Fujifilm WAKO 019-19741, 1:500), rabbit-anti-P2Y12 (Invitrogen 702516; 1:500), rat-anti-CD68 (Bio-rad MCA1957T; 1:400), chicken-anti-RFP (Rockland Immunochemicals 600-901-379; 1:1,000), chicken-anti-GFAP (Abcam AB4674; 1:1,000), rabbit-anti-S100β (Abcam AB41548; 1:500), and rabbit-anti-Olig2 (Merck AB9610; 1:500). After washing in PBS and 0.3% Triton-X100 (PBS-T; 3 × 20 min), incubation with secondary antibody (in PBS-T with 3% BSA) was carried out for 24 h on a nutator at 4°C, with the following antibodies: goat-anti-rabbit-568 (Thermo Fisher A11011; 1:1,000), goat-anti-rabbit-647 (Thermo Fisher A21244; 1:1,000), goat-anti-chicken-488 (Thermo Fisher A11039; 1:1,000), goat-anti-chicken-568 (Thermo Fisher A11008: 1:1000), and goat-anti-rat-488 (Thermo Fisher A11006; 1:1,000). After 5 × 20 min washes in PBS-T, stained slices were mounted to Superfrost Plus slides (VWR, 48311-703) with DAPI-containing Vectashield mounting medium (Vector Laboratories, H-1200)

#### c-Fos Immunostaining

Following slice collection (60 µm), slices were first incubated in 4% PFA for 10 min. Then, slices were incubated in blocking buffer for 3 h at room temperature before being washed 3 times with PBS. Slices were incubated with rabbit-anti-cFOS (1:1,000, Sysy Antibodies, 226 008) in PBS overnight at 4°C on a nutator. Slices were next washed 5 times with PBS with 0.3% Triton-X100 (PBST) before a secondary incubation with goat-anti-rabbit-647 (Thermo Fisher A21244; 1:1,500) overnight at 4°C on a nutator. Slices were then washed 5 times with PBST, and 3 times with PBS before mounting to Superfrost Plus slides (VWR, 48311-703) with DAPI-containing Vectashield mounting medium (Vector Laboratories, H-1200).

#### In situ hybridization and immunostaining

Mice were transcardially perfused with 20 mL phosphate-buffered saline (PBS) and brains dissected and embedded in OCT (Tissue-Tek, 4583), frozen on dry ice and stored at –80°C. 18 µm cryosections through the MPOA were taken using a cryostat (Leica CM1950) and collected on Superfrost Plus slides (VWR, 48311-703). Slides were fixed in 10% Neutral buffered formalin (NBF), followed by a series of dehydration steps in ethanol (5 min each of 50%, 70%, 100%, 100% v/v EtOH). Slides were pre-treated with RNAscope Protease III reagent. Single-molecule fluorescent *in situ* hybridisation was performed on slides using the RNAscope LS Multiplex Reagent Kit (Advanced Cell Diagnostics), LS 4-Plex Ancillary Kit and Multiplex Reagent Kit on a robotic staining system (Leica BOND-III). RNAscope probes were *Gal* (Cat. No. 400968), *Fos* (Cat. No. 316928), *Esr1* (Cat. No. 478208), *Pgr* (Cat. No. 318928), *Prlr* (Cat. No. 430791), *Aldh1L1* (Cat. No. 405898), and *Slc1a3* (Cat. No. 430788).

NeuN antibody (Millipore MAB377) was used at 1:500 after completing *in situ* hybridisations.

### Surgical procedures and viral injections

#### Surgeries on neonates (P0–1)

The entire litter including nest was removed from the home cage and kept in a heated recovery chamber for the duration of the surgeries. Mice were individually removed from the chamber and anaesthetised with isoflurane (5% induction, 4% maintenance in oxygen-enriched air). P0/1 injections were carried out on a modified stereotaxic frame (Model 940, Kopf Instruments) where they were head fixed and isoflurane was delivered through a 3D printed nose cone^65^. An incision in the scalp was made with fine scissors, and coordinates were determined from lambda. A 25G needle was then used to gently incise the skull. Viruses were injected using a Nanoject II or Nanoject III injector (Drummond Scientific) with pulled glass capillary tubes (3-000-203-G/X, Drummond Scientific). The needle was left *in situ* for 5–7 min and then slowly removed. Surgery sites were closed with Vicryl sutures (Ethicon) and Vetbond (3M). Mice were returned to the heated recovery chamber after surgery. After all surgeries on the litter were completed, pups were returned to their home cage along with the nest and left undisturbed for at least 5 days.

#### Surgeries on juveniles (≥ P10)

An entire litter of mice pre-weaning were removed from the home cage with nest and kept in a heated recovery chamber for the duration of the surgeries. Mice were individually removed from the chamber for surgeries. Prior to surgery, mice were given meloxicam (10 mg/kg) orally before surgery. Adults were individually removed from their home cages for surgery. Mice were anaesthetised with isoflurane (5% induction, 1.5% maintenance in oxygen-enriched air) and head-fixed on a stereotaxic frame (Model 940, Kopf Instruments).

Meloxicam (10 mg/kg) and buprenorphine (0.1 mg/kg) were given subcutaneously prior to surgery and eyes were projected throughout using ophthalmic ointment (Viscotears, Alcon). Rectal body temperature was maintained throughout surgery at 37°C using a heating pad (Havard Apparatus). A hole was drilled in the skull and viruses were injected using a Nanoject II or Nanoject III injector (Drummond Scientific) with pulled glass capillary tubes (3-000-203-G/X, Drummond Scientific). The needle was left *in situ* for 5–7 min and then slowly removed. Surgery sites were closed with Vicryl sutures (Ethicon) and Vetbond (3M). Mice were returned to the heated recovery chamber, post-surgery; pre-weaned mice were returned all together post-surgery along with the nest. Adults were returned to the home cage once mobile. Adults were provided with oral meloxicam (10 mg/kg) for 3 days post-surgery.

#### Viral injections

For anterograde tracing experiments, P0–1 *Gal-Cre* mice were unilaterally injected with 60 nL of a 1:1 mix of AAV2/1-CAG-FLEx-tdTomato (Addgene 28306, 2.5 × 10^13^ GC/mL) and AAV2/1-CAG-FLEx-SynGFP (gift from Silvia Arber)^66^ into the MPOA. Adult *Gal-Cre* mice were injected with 250 nL of the above 1:1 virus mix into the MPOA. For chemogenetic experiments, neonate (P0–1) *Gal-Cre* mice were unilaterally injected with 60 nL of AAV2/1-CAG-FLEx-hM4Di-mCherry (Crick Vector Core, 1.5 × 10^12^ GC/mL) or 60 nL AAV2/1-CAG-FLEx-hM3Dq-mCherry (Crick Vector Core, 7.8 × 10^12^ GC/mL). MPOA coordinates were from lambda in juveniles (AP: +1.93 mm; ML: +0.15 mm; DV: –3.75 mm) and from bregma in adults (AP: 0 mm; ML: +0.5 mm; DV: –5.05 mm).

For retrograde tracing experiments, neonate (P0–1) *Gal-Cre* mice were unilaterally injected with 60 nL of a 1:1 mix of starter viruses AAV2/1-Syn-FLEx-nEBFP-E2A-TVA (Crick Vector Core, 5.4 × 10^13^ GC/mL) and AAV2/1-Syn-FLEx-nCherry-N2cG (Crick Vector Core, 6.9 × 10^13^ GC/mL) at the MPOA with the above coordinates. At either P10 or P15, mice were injected with 300 nL of G-deleted rabies virus (EnvA-RΔG-B19-EGFP, Crick Vector Core, 5.8 × 10^8^ GC/mL) into the MPOA. Coordinates from bregma were AP: 0 mm; ML: 0.3 mm; DV: – 4.75 mm. Adult *Gal-Cre* mice were injected with 250 nL of the 1:1 starter mix into the MPOA. 10 days later, they were injected again at the MPOA with 300 nL of the G-deleted rabies virus. Confirmatory anterograde tracing of the retrograde tracing was carried out from area CA1–3 of the hippocampus. C57BL/6J mice were injected with a 1:1 mix of AAV2/1-CAG-tdTomato (Addgene 28306, 2.5 × 10^13^ GC/mL) and AAV2/1-CAG-SynGFP (Crick Vector Core, 3.12 × 10^12^ GC/mL). P0 mice were injected with 60 nL at coordinates AP: 1.45 mm; ML: 0.6 mm; DV: –1.1 mm from lambda. Adults were injected with 250 nL at coordinates AP: –1.6 mm; ML: 1.2 mm; DV: –1.52 mm from bregma.

#### Ovariectomy

Entire litters of P7 mice were removed from the home cage along with the nest and placed in a heated recovery chamber for the duration of the surgeries. Prior to induction, mice received oral meloxicam (10 mg/kg). Anaesthesia was provided via isoflurane (5% induction, 1.5% maintenance) in oxygen-enriched air (1 L/min), and mice were secured in a stereotaxic frame (Model 940, Kopf Instruments) via a custom nose cone. Meloxicam (10 mg/kg) and buprenorphine (0.1 mg/kg) were administered subcutaneously. A small dorsal incision was made through skin and muscle, and the ovaries were bilaterally exteriorised and removed. Incisions were closed using Vetbond (3M), and mice were returned to the heated chamber before reuniting the entire litter and nest with the dam. Oral meloxicam (10 mg/kg) was continued for three days post-operatively.

### Fibre photometry

Neonate (P0–1) *Gal-Cre* mice were injected with 200 nL of AAV2/1-CAG-FLEx-GCaMP8s (Addgene 162380, 1.9 × 10^13^ GC/mL) at the MPOA. At P9–10, mice were implanted with a unilateral fibre-optic cannula (MFC_200/240-0.22_6.0mm_MF1.25_FLT, Doric Lenses) above the injected MPOA hemisphere at AP: 0 mm ML: +0.3 mm DV: –4.75 mm from bregma. Implants were fixed to the skull with UV light-curable glue (RelyX, Unicem, 3M) and Jet acrylic resin (Lang Dental). Incisions were closed over the cement using Vicryl sutures (Ethicon) and Vetbond (3M). Set-up in terms of experimental timing and testing was performed similar to the pup-directed behaviour assay. Mice were connected to a custom patch-cord (Doric Lenses) as part of a FP3001 fibre photometry system (Neurophotometrics). Briefly, two LEDs (415 nm, 470 nm, light power ∼50 µW) were pulsed at 20 Hz in an interleaved manner to obtain an isosbestic motion signal (415 nm) and GCaMP fluorescence (470 nm). A FLIR 277 BlackFly CMOS camera was used to detect fluorescent signals, and acquisition was controlled (and synchronised to the acquisition of behavioural video recordings) via Bonsai (https://bonsai-rx.org/). Mice were presented with salient stimuli in a random order during the trial, ending with mouse milk pipetted into the mouse. Mouse milk was collected using a 10cc syringe from lactating dams following oxytocin injection (100 IU, SigmaAldrich, O4375). Z-scores were calculated between –3 and –1 s prior to (baseline), and 1–3 s after (response), initial contact with the stimulus. Distributions of averaged, Z-scored responses for each mouse at each postnatal day were compared to the distributions of their corresponding baseline periods using a Wilcoxon signed-rank test with Benjamini-Hochberg FDR.

### Imaging

#### Wide-field imaging

Wide-field imaging of tissue was performed using a Vectra Polaris Automated Quantitative Pathology Imaging System (Akoya Biosciences) at 20× magnification. Using custom QuPath^67^ scripts, fluorescent cell bodies were detected and colocalised where appropriate.

#### Serial two-photon tomography

Rabies-injected brains were imbedded in 4% agarose in 0.1 M phosphate buffer and imaged using serial-section two-photon tomography^68,69^. 8 serial sections were imaged each 5 µm with 2 µm by 2 µm lateral resolution, after which a 40 µm physical section was cut by vibrating blade. Images were taken at 920 nm and filtered for the primary EGFP signal, and a far-red background signal. Individual images were stitched using the MATLAB plugin *StitchIt* (https://github.com/SWC-Advanced-Microscopy/StitchIt) to form a full coronal section. Brains were registered to the Allen Mouse Brain CCFv3^70^ using the open-source *brainreg* package^71^, and rabies expressing neurons were detected in 3D and curated with the *cellfinder* package^72^. Detected cells coordinates were downsampled to 10 µm to match the Allen Mouse Brain CCFv3 space. Convergence index was calculated as the number of rabies expressing neurons per brain region divided by the number of starter cells. Input fraction was calculated as the number of rabies expressing neurons per brain region divided by the total number of rabies expressing neurons. Fold changes were calculated by dividing mean values of later time points (e.g., adult) by mean values of earlier time points (e.g., P14).

### Slice electrophysiology

*Gal-Cre; tdTomato* mice were used for targeted recordings of MPOA^Gal^ neurons. Mice were rapidly decapitated, and the brain was quickly dissected out into an ice-cold slicing solution containing (in mM): sucrose (214), KCl (2), NaH_2_PO_4_ (1.2), NaHCO_3_ (26), MgCl_2_ (2), CaCl_2_ (2), and D-glucose (10), equilibrated with carbogen (95% O_2_ / 5% CO_2_). Acute 200 µm coronal brain slices containing the MPOA were taken using a Vibratome (Leica VT1200S) and incubated at 35°C in a chamber of artificial cerebrospinal fluid (aCSF) containing (in mM): NaCl (127), KCl (2), NaH_2_PO_4_ (1.2), NaHCO_3_ (26), MgCl_2_ (1.3), CaCl_2_ (2.4) and D-glucose (10) continually perfused with carbogen. After at least 1 h of recovery, slices were transferred to a recording chamber under an upright microscope with infrared Nomarski differential interference contrast optics (Slicescope, Scientifica). Slices were superfused with carbogenated aCSF (1–2 mL/min) at ∼33°C during experiments. Glass micropipettes (3–6 MΩ resistance) were pulled from borosilicate capillaries (World Precision Instruments, Aston, UK) on a P-97 Flaming/Brown micropipette puller (Sutter, Novato, CA) and filled with internal solution containing (in mM): K-Gluconate (140), KCl (10), KOH (1), EGTA (1), Na_2_-ATP (2), HEPES (10), pH 7.3, 280–290 mOsm. Voltage clamp recordings were performed at a holding potential of −60 mV. For *post hoc* reconstruction of neuronal morphology, neurobiotin (0.2% [w/v], SP-1120, Vector Laboratories) was added. Following whole-cell access to tdTomato^+^ neurons, access resistance was monitored throughout the experiment, and neurons in which the series resistance exceeded 25 MΩ or changed by > 20% were excluded. Resting membrane potential was assessed immediately after establishing a whole-cell configuration. Recordings were performed using an Axon Instruments Multiclamp 700B amplifier (Molecular Devices), low pass filtered at 10 kHz and digitized using a Digidata 1550B digitizer (Molecular Devices). Slow and fast capacitive components were semi-automatically compensated. Offline data analysis was performed with Clampfit 10 software (Molecular Devices) and WinEDR v4 (http://spider.science.strath.ac.uk/sipbs/software_ses.htm).

### Morphological reconstructions

#### Neuronal reconstructions

One tdTomato^+^ neuron per MPOA hemisphere (200 µm thick slices) was recorded and filled with 2% neurobiotin. Following recordings, slices were post-fixed with 4% PFA in PBS with sucrose (200 mM) and HEPES (100 mM) at 4°C overnight on a nutator, rinsed in PBS and washed in PBS-T for 1 h. Incubation with streptavidin-633 (ThermoFisher S21375, 1:1,000) was performed for 24 h at 4°C on a nutator. After washing in PBS-T for 1 h, slices were rinsed in PBS and mounted on slides in DAPI-containing Vectashield mounting medium (Vector Laboratories, H-1200). Brain slices were imaged on an inverted Olympus CSU-W1 SoRa confocal microscope, using a 60× (NA 1.4) oil immersion objective and a Z step size of 0.5 µm. Morphological reconstruction of the soma, projections and spines of each neurobiotin-filled neuron was performed manually using the Filament Tracer module (auto-path method, spine diameter 0.25 µm) in Imaris 9.0 (Bitplane). Axons were considered as those projections which do not have any spines along their length. Ramification index was calculated as the ratio of endpoints to neurites. Ramification fraction was calculated as the ratio of endpoints per dendrite or axon to the total neurite number. To assess neuronal arbour complexity, Sholl analysis on reconstructed microglia was performed in Imaris by quantifying intersections with concentric spheres (1 µm interval) centred on the soma.

#### Glial reconstructions

Image volumes (0.5 µm steps, 60× oil-immersion magnification) of IBA1-stained regions in MPOA, MOp/s and CPu were acquired using an inverted Olympus CSU-W1 SoRa confocal microscope. IMARIS (Version 10.1; Oxford Instruments) was used to surface render IBA1 staining with 0.1 µm smoothing following the protocol by Nemes-Baran & DeSilva (2021)^73^. Microglia that were only partially imaged were removed from analysis. Small surfaces of either unconnected IBA1 staining or bright background were manually removed. Fused microglia were manually separated using the cut function. The *mask all* function was used on the resulting surfaces to create the final surface reconstruction for skeletonisation of the remaining IBA1 signal. Skeletonisation of microglia was carried out following Althammer *et al.,* (2020)^74^ using the filament function with the following settings: detect new starting points: largest diameter 3.95 µm, seed points 0.3 µm; remove seed points around starting points: diameter of sphere regions: 15 µm. Seed points were either placed or manually removed from the soma if they were incorrectly identified. All surface and filament parameters were exported to individual .csv files for analysis. To assess microglial complexity, Sholl analysis on reconstructed microglia was performed in Imaris by quantifying intersections with concentric spheres (1 µm interval) centred on the soma.

### Microglia depletion

Juvenile mice were injected intraperitoneally with 100 mg/kg of BLZ945 (MedChem Express, HY-12768) dissolved in 20% [w/v] (2-Hydroxypropyl)-β-cyclodextrin (Sigma Aldrich, H107) in distilled water. Injections were delivered daily for 4 days prior to behavioural testing or electrophysiology.

### Quantifications and statistical analyses

Details of statistical tests are included in figure legends. Statistics and *P* values were calculated using *scipy* stats module (1.13.0). Parametricity was determined by Shapiro-Wilk test. Graphs were produced using GraphPad Prism (10.4.2). Significance levels are: \*\*\**P* < 0.001; \*\**P* < 0.01; \**P* < 0.05.

#### Image analysis and registration

The Fiji plugin ABBA^75^ was used to register coronal brain sections to the Allen Brain Atlas (CCFv3)^70^. Briefly, X and Y rotation were adjusted across all sections from a given brain and underwent non-rigid registration using the BigWarp tool (sample channel: DAPI, atlas channel: Nissl). Positive cell detection was performed on the transformed samples in QuPath^67^.

#### Enrichment ratios

Enrichment was calculated as the proportion of *c-fos⁺* cells co-expressing a specific marker (e.g., *Gal*), divided by the proportion of *c-fos⁺* cells co-expressing NeuN. NeuN serves as a reference for baseline expression, yielding an enrichment ratio of 1 when the marker of interest is expressed at the same frequency as NeuN.

#### Specificity index

To quantify tuning specificity, a tuning specificity index (TSI) was computed from Z-scored fibre photometry signals. Z-scores were first calculated by subtracting baseline fluorescence from stimulus-evoked responses and then averaged across stimulus presentations for each condition. For each stimulus, the mean response (*x̄_s_*) was compared against the mean of the pooled responses to all other stimuli (*x̄_r_*) and normalised by the pooled standard deviation (*σ)* to account for response variability. The TSI is defined as:

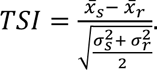

To assess statistical significance, non-parametric bootstrapping (n = 1,000 iterations) was performed using *scikit-learn* v1.2.2. The proportion of bootstrap samples with TSI > 0 was used to estimate a one-sided *P* value. TSI > 0 indicates positive tuning, reflecting stronger responses to the stimulus of interest (e.g., pups) than to other stimuli. A TSI ≈ 0 indicates no tuning, with comparable responses across stimuli. A TSI < 0 indicates negative tuning or suppression, i.e., responses to the stimulus of interest that are weaker than to other stimuli.

#### Unsupervised clustering

Unsupervised clustering was carried out on dendritic spines based on 13 morphological parameters (Extended Data Fig. 12) and in microglia based on 8 morphological parameters (Extended Data Fig. 14). Using *scikit-learn* v1.2.2 data were standardised using Z-score normalisation before K-means clustering into 4 groups. The number of groups was determined via the elbow point method prior to clustering.

#### Dimensionality reduction

Dimensionality reduction was performed via Principal Component Analysis (PCA) using *scikit-learn* v1.2.2. For anterograde and retrograde tracing data, predefined groups were assigned based on animal age. Anterograde parameters were based on the density of fibres and presynaptic puncta in each brain region. Data were normalised to MPOA values to account for variations in viral transduction. Retrograde parameters were based on the convergence index of each brain region. For dendritic spines and microglia, K-means clustering (see ‘Unsupervised clustering’) was performed prior to dimensionality reduction.

To visualise sample distribution and group separation, data was plotted in 2D retaining the first two PCs. Feature contributions for the first two PCs were extracted from the data to determine the relative influence of each variable on the reduced dimensions. Pairwise Euclidean distances were calculated using *scipy* (1.11.4) to assess inter-cluster relationships.

## Acknowledgments

We thank A. Schaefer, F. Iacaruso, S. Ultanir, L. Arancibia Carcamo and members of the State-dependent Neural Processing Lab for discussions and comments on the manuscript. We thank the Biological Research and Surgical Services Facilities at the Francis Crick Institute for animal care and technical assistance, as well as Crick Light Microscopy, Experimental Histopathology, Making Lab, Mechanical Workshop, and Vector Core. We thank M.-J. Dolan for advice on microglia classification and manipulation. This study received support from the Francis Crick Institute, core funding FC001153. The Francis Crick Institute receives its funding from Cancer Research UK, the UK Medical Research Council, and the Wellcome Trust (J.K.). This study was also supported by European Research Council starting grant ERC-2019-STG847873 (J.K.) and a British Society for Neuroendocrinology Project Support Grant (B.B.J.).

## Author contributions

B.B.J. and J.K. conceptualised the study. B.B.J., M.X.C. and J.K. contributed to the methodology. B.B.J., M.X.C., S.L., L.S.H.E.R., and G.M.K.C. performed the investigation. B.B.J. and J.K. performed the formal analysis. B.B.J. and J.K. acquired funding. B.B.J. and J.K. supervised the project. B.B.J. and J.K. wrote the original draft of the manuscript. B.B.J. and J.K. reviewed and edited the manuscript.

